# Iron-sulfur clusters are involved in post-translational arginylation

**DOI:** 10.1101/2021.04.13.439645

**Authors:** Verna Van, Janae B. Brown, Hannah Rosenbach, Ijaz Mohamed, Nna-Emeka Ejimogu, Toan S. Bui, Veronika A. Szalai, Kelly N. Chacón, Ingrid Span, Aaron T. Smith

## Abstract

Eukaryotic arginylation is an essential post-translational modification that both modulates protein stability and regulates protein half-life through the N-degron pathway. Arginylation is catalyzed by a family of enzymes known as the arginyl-tRNA transferases (ATE1s), which are conserved across the eukaryotic domain. Despite its conservation and importance, little is known regarding the structure, mechanism, and regulation of ATE1s. In this work, we have discovered that ATE1s bind a previously unknown iron-sulfur cluster that is conserved across evolution. We have extensively characterized the nature of this iron-sulfur cluster, and we show that the presence of the iron-sulfur cluster is linked to alterations in arginylation efficacy. Finally, we demonstrate that the ATE1 iron-sulfur cluster is oxygen sensitive, which could be a molecular mechanism of the N-degron pathway to sense oxidative stress. Thus, our data provide the framework of a cluster-based paradigm of ATE1 regulatory control.

## INTRODUCTION

In eukaryotes, the half-life of a protein is largely regulated by the ubiquitin-proteasome system (UPS), which catalyzes the covalent attachment of the small protein ubiquitin (Ub) to Lys residues of polypeptides, providing a molecular flag for degradation via the 26S proteasome.^1–4^ The generation of degradation signals is dependent on the presence of an internal Lys residue (the site of the formation of the poly-Ub chain) and the identity of the N-terminal residue, the latter giving rise to the aptly named N-degron pathway (formerly known as the N-end rule pathway).^5–7^ A major component of this pathway is the three-level hierarchical Arg N-degron branch that requires downstream processing prior to Ub conjugation (Fig. 1a). As the first step of the Arg N-degron pathway, tertiary destabilizing N-terminal residues Asn, Gln, and Cys require modification by specific enzymes, dioxygen (O_2_), or nitric oxide (NO), before further recognition. Specifically, Asn and Gln must be deamidated to Asp and Glu by the respective enzymes N-terminal asparagine amidohydrolase (NTAN) and N-terminal glutamine amidohydrolase (NTAQ),^8^ whereas Cys must be oxidized to Cys-sulfinic or sulfonic acid.^9^ These modifications yield secondary destabilizing residues, which are subsequently recognized by arginine transferases (known as ATE1s), essential enzymes of the Arg N-degron pathway that arginylate N-terminal residues in a tRNA-dependent manner to form primary destabilizing residues (Fig. 1a).^10^ Finally, arginylated proteins may be recognized by E3 ubiquitin ligases, ubiquitinated, and transported to the proteasome for degradation (Fig. 1b), or they may exhibit non-degradative alteration in function (Fig. 1c).^6,11–13^

**Figure 1.**
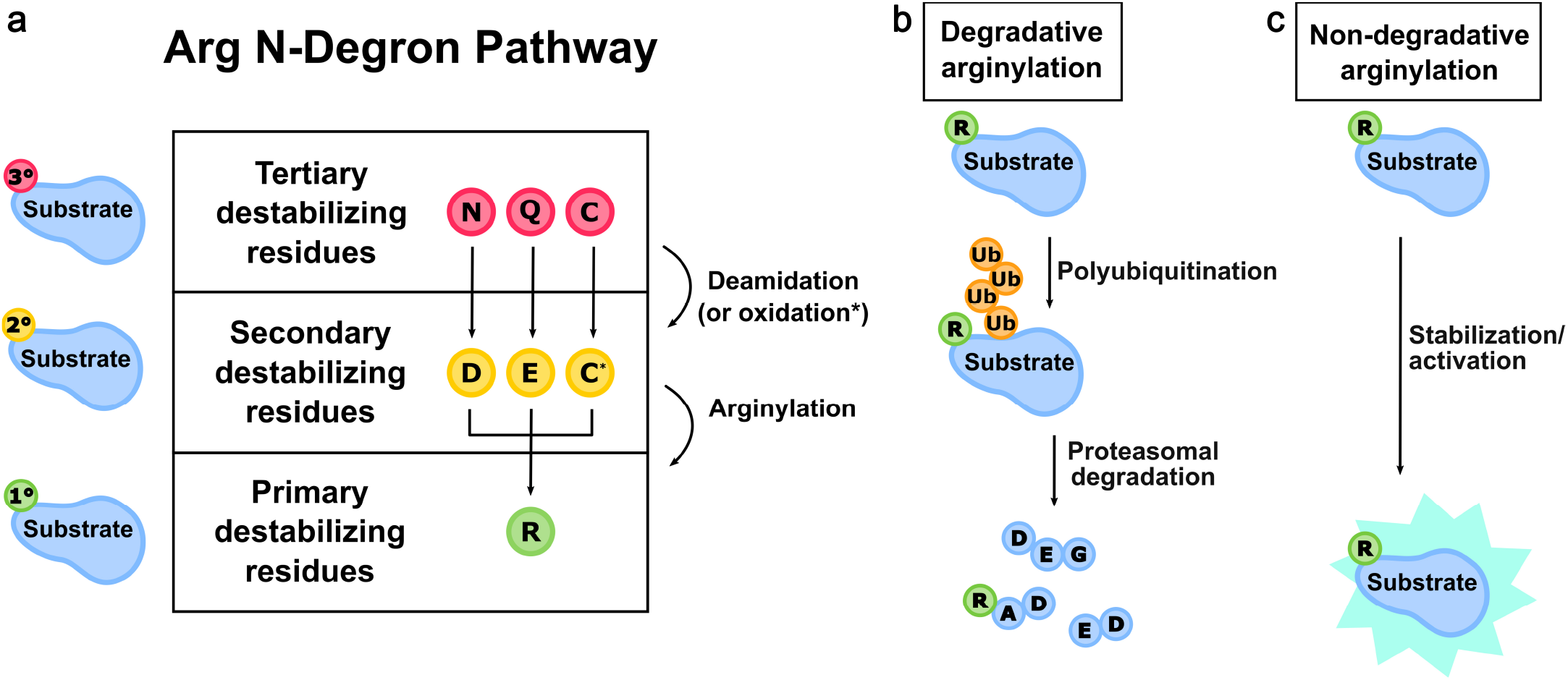
The effects of post-translational arginylation. **a**. The eukaryotic Arg N-degron pathway is a hierarchical pathway that controls the fate of many proteins in the cell. Tertiary destabilizing residues (red circles) are N-terminal Asn, Gln, and/or Cys residues that are exposed upon protein processing. These residues are modified via deamidation or oxidation to form the secondary destabilizing residues (yellow circles) Asp, Glu, and/or oxidized Cys (C*). Secondary destabilizing residues are recognized by arginine transferases (ATE1s), which enzymatically transfer Arg to the N-terminus of a polypeptide, resulting in the exposure of a primary destabilizing residue (green circle). **b**. Via the degradative pathway, arginylated substrate polypeptides are recognized by N-recognins, polyubiquitinated, and broken down in a proteasomal manner. **c**. Via the non-degradative pathway, arginylated substrate polypeptides may have altered stability, oligomerization, and even activity.

The fidelity of the Arg N-degron pathway is critical for normal eukaryotic cellular function.^4^ For example, key protein targets of this pathway are responsible for several essential physiological processes from chromosomal segregation to the stress response.^7,14–17^ Additionally, important studies have demonstrated a link between ATE1 function, the sensing of O_2_ availability via plant cysteine dioxygenases (CDOs), and the oxidized Cys component of the Arg N-degron pathway.^18–20^ Further underscoring the importance of this post-translational modification, the Arg N-degron branch also plays an essential role in mammalian cardiovascular maturation and blood pressure control. Data link this pathway and ATE1s directly to cardiovascular development,^9,16,21,22^ to controlling signaling of mammalian G proteins that regulate Ca^2+^ homeostasis,^23^ and to thrombosis as well as progressive heart failure in murine models lacking ATE1,^24^ highlighting the significance of arginylation in vascular health and disease. However, target proteins of ATE1-catalyzed arginylation are highly diverse and have varied N-terminal residues, requiring regulation at multiple levels for faithful arginylation and protein turnover.

Regulation of ATE1 may occur at the post-transcriptional as well as post-translational levels.^4^ In some lower-order eukaryotes such as single-celled yeasts, only one (iso)form is thought to exist encoded along a single gene^25^, while higher-order eukaryotes may have more than one *ate1* gene that can encode for several isoforms.^12,26,27^ Once translated, these different isoforms may differ by as little as *ca*. 5000 g/mol (5 kDa).^10,28^ However, as has been reported for *Mus musculus* (mouse) ATE1, these isoforms may exhibit important differences in tissue distribution, protein specificity, and even arginylation efficacy,^10,29–31^ demonstrating post-transcriptional regulation of the *ate1* gene. At the post-translational level, the ATE1-controlled Arg N-degron pathway may be regulated by the presence of small molecules such as O_2_, NO, and iron protoporphyrin IX (heme *b*).^19,21,32–34^ These results tantalizingly suggested that ATE1s could be heme-binding proteins that respond to small molecules in a manner similar to well-known gas-sensing proteins.^35,36^ However, the precise molecular-level details of this regulatory process are essentially unknown.

To begin our investigations, we sought to probe the heme-binding properties of yeast (*Saccharomyces cerevisiae*) ATE1 (*Sc*ATE1). In a surprising turn of events, we found that *Sc*ATE1 binds an [Fe-S] cluster. Using a mixture of spectroscopic, biochemical, and biophysical techniques, we probed the as-isolated and reconstituted cluster-binding environments of *Sc*ATE1. These data demonstrate that *Sc*ATE1 binds a [4Fe-4S]^2+/+^ cluster in its N-terminal domain that is oxygen-sensitive, redox active, and capable of being converted to a [2Fe-2S]^2+^ cluster under oxic conditions. Importantly, we reveal that the presence of the [Fe-S] cluster can affect the *in vitro* arginylation efficacy of *Sc*ATE1. Lastly, using bioinformatics data, expression of mouse ATE1 isoform 1B7A, and reconstitution approaches, we show that the presence of an [Fe-S] cluster is an evolutionarily conserved attribute of ATE1s. Based on these results, we propose an [Fe-S] cluster-based paradigm of ATE1 regulatory control, which may be operative within the Arg N-degron pathway across eukaryotes.

## RESULTS

### *Sc*ATE1 unexpectedly binds an [Fe-S] cluster

We initially set out to characterize the heme-binding properties of yeast (*Saccharomyces cerevisiae*) ATE1 (*Sc*ATE1), which exists as a single, soluble isoform. We cloned, expressed in *Escherichia coli*, and purified His-tagged *Sc*ATE1 (518 amino acids and 59000 g/mol *i.e.*, 59.6 kDa; Supplementary Figure 1a, verified by Western blotting) to high purity and high oligomeric homogeneity (chiefly monomeric) as judged by SDS-PAGE and gel filtration (Supplementary Figure 1b,c). Importantly, concentrated *Sc*ATE1 was a slightly brown color, indicating the presence of a putative cofactor, which we initially suspected was heme. However, metal analyses indicated very little iron (0.07 Fe ion ± 0.04 Fe ion per polypeptide, *n* = 5), and the electronic absorption (EA) spectrum of this protein indicated no spectral features consistent with the presence of heme. Initially surmising that *Sc*ATE1 was a hemoprotein, and suspecting poor cofactor incorporation during expression, we supplemented our growth media with known heme *b* precursors such as ferric (Fe^3+^) citrate and δ-aminolevulinic acid (ALA), which is a common strategy for increasing accumulation of heme-replete proteins.^37,38^ Despite a modest color change in our cell pellets, and a slightly darker brown color of purified *Sc*ATE1 (Supplementary Figure 1d, inset), the analyzed spectral features (Supplementary Figure 1d) were wholly inconsistent with the presence of heme *b* in the purified enzyme.^39,40^ They were, instead, remarkably similar to proteins containing iron-sulfur ([Fe-S]) clusters.^41,42^

Presuming that *Sc*ATE1 might actually be an [Fe-S] cluster-binding protein, we modified our expression and purification protocol to facilitate cluster incorporation in several ways: media supplementation with ferric citrate and *L-*Cys, as well as co-expression with a high-copy plasmid encoding the *E. coli* iron-sulfur cluster (ISC) biosynthesis machinery (pACYC184*iscS-fdx*).^43^ Although expression of *Sc*ATE1 in the presence of only ferric citrate and *L-*Cys dramatically changed the color of the *E. coli* culture and appeared to increase substantively the spectral features associated with the cluster, reproducibility was problematic. We finally achieved high reproducibility of *Sc*ATE1 expression in the presence of pACYC184*iscS-fdx* (constitutively active at low levels) in media supplemented with ferric citrate and *L-*Cys. After expression under these conditions and oxic purification, the darkly colored *Sc*ATE1 (Fig. 2a), which maintained its monomeric oligomerization, exhibited an electronic absorption (EA) spectrum diagnostic of a [2Fe-2S]^2+^ cluster (Fig. 2b).

**Fig. 2.**
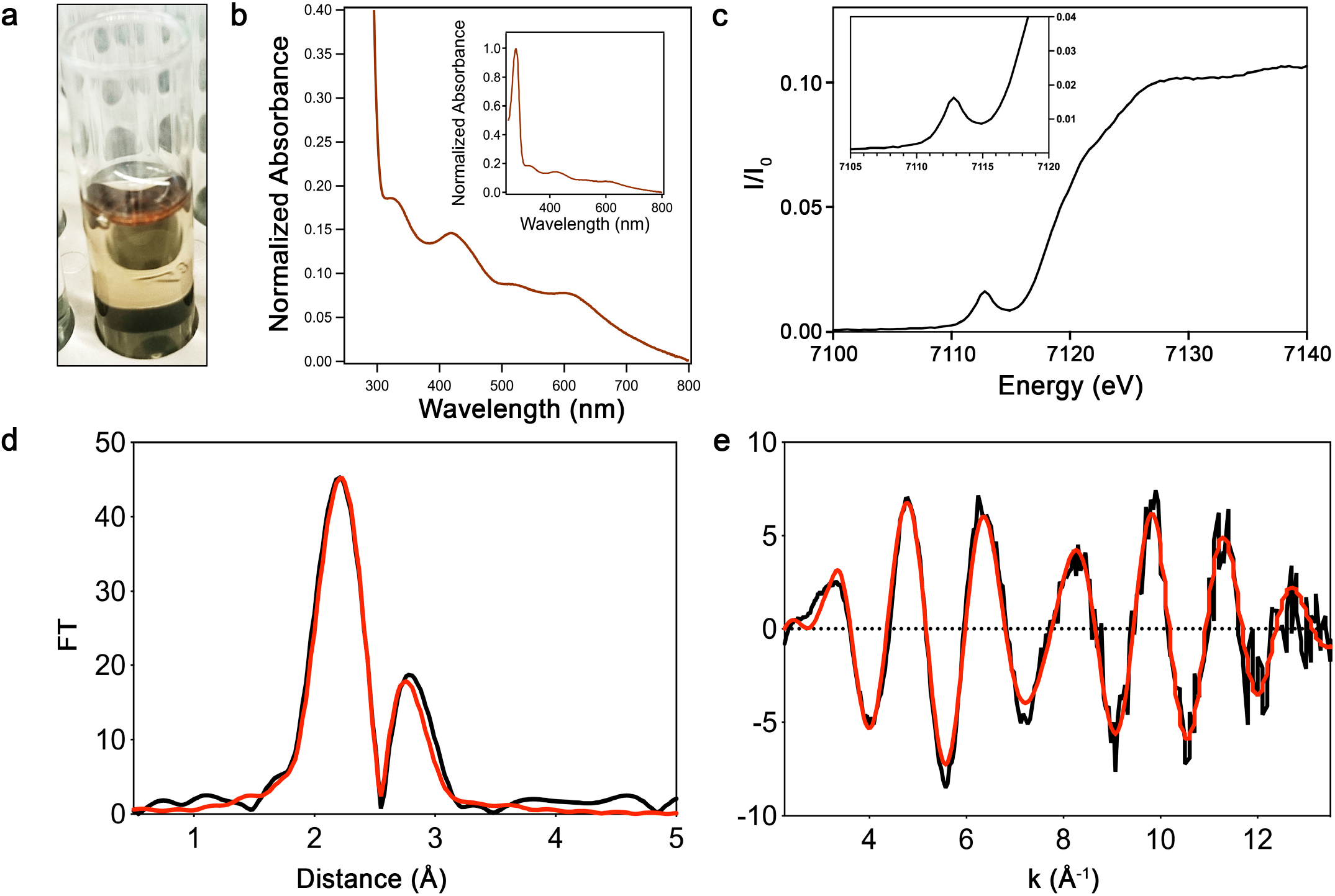
*Sc*ATE1 binds a [2Fe-2S]2+ cluster when purified under oxic conditions. **a**. Purified *Sc*ATE1 has a brown color, consistent with the presence of an [Fe-S] cluster. **b**. Features of the visible electronic absorption spectrum of *Sc*ATE1 isolated under oxic conditions is consistent with the presence of a [2Fe-2S]2+ cluster. Inset: expanded absorption spectrum showing both UV and visible regions. **c**. The Fe XANES of *Sc*ATE1 isolated under oxic conditions. Inset: expanded pre-edge feature of the 1s→3d transition at 7112.5 eV, consistent with tetrahedral Fe3+ sites. **d**,**e**. EXAFS of *Sc*ATE1 isolated under oxic conditions. The Fourier transformed data (**d**) and raw EXAFS (**e**) are consistent with the presence of a [2Fe-2S]2+ cluster. 1 Å = 0.1 nm.

Characterization of *Sc*ATE1 expressed in the presence of the ISC machinery and isolated under an oxic atmosphere is most consistent with a single ferredoxin-like [2Fe-2S]^2+^ cluster bound to the polypeptide. Analysis of acid-labile iron using the ferrozine assay indicated the presence of (1.21 ± 0.26) mole eq. of iron per polypeptide (*n* = 3). This analysis is consistent with the [2Fe-2S] designation, assuming the production of some apo protein, which is not uncommon for cofactor incorporation during heterologous expression. The [2Fe-2S] assignment is confirmed based on the features of the EA spectrum of *Sc*ATE1 isolated under oxic conditions (Fig. 2b) and further spectroscopic characterization (*vide infra*). The *Sc*ATE1 EA spectrum of protein purified under oxic conditions displays characteristic absorption maxima in the near-UV and visible regions: *ca*. 330 nm (shoulder), 414 nm, *ca*. 475 nm (shoulder), and a broad absorbance envelope from 500 nm to 700 nm, similar to proteins bearing [2Fe-2S]^2+^ ferredoxin-like clusters.^43–47^ This absorbance spectrum is accompanied by a weak circular dichroism (CD) spectrum in the visible region (Supplementary Figure 2), indicating the presence of the cluster within the polypeptide.

To characterize the [Fe-S] cluster further, we analyzed *Sc*ATE1 expressed in the presence of the ISC machinery and isolated under oxic conditions using X-ray absorption spectroscopy (XAS). The X-ray absorption near edge structure (XANES) is shown in Fig. 2c and displays a distinct pre-edge feature at 7112.5 eV, which is characteristic of tetrahedral Fe^3+^ centers. Further evidence for this oxidation state assignment is the first inflection point of the edge at ≈ 7120 eV, common for Fe^3+^ centers (Fig. 2c). Simulations of the extended X-ray absorption fine structure (EXAFS) data of *Sc*ATE1 reveal only S-based environments as the nearest neighbor ligands with an Fe-S distance of 2.25 Å (1 Å = 0.1 nm; Fig. 2d,e; Table 1). Furthermore, there is no indication of O/N-nearest neighbor ligands, nor any indication of Fe-C scattering, precluding the involvement of the (His)_6_ tag in iron binding, and ruling out the Rieske-(2 Cys/2 His) or mitoNEET-type (3 Cys/1 His) [2Fe-2S] cluster types. Long-range scattering interactions representing only a single Fe-Fe vector are observed and fitted to a distance of 2.71 Å (Fig. 2d; Table 1), strongly supporting the [2Fe-2S] designation. Altogether, the metal analyses, EA, and XAS data clearly point to the presence of a single ferredoxin-like [2Fe-2S]^2+^ cluster in oxically-purified *Sc*ATE1.

**Table 1.**
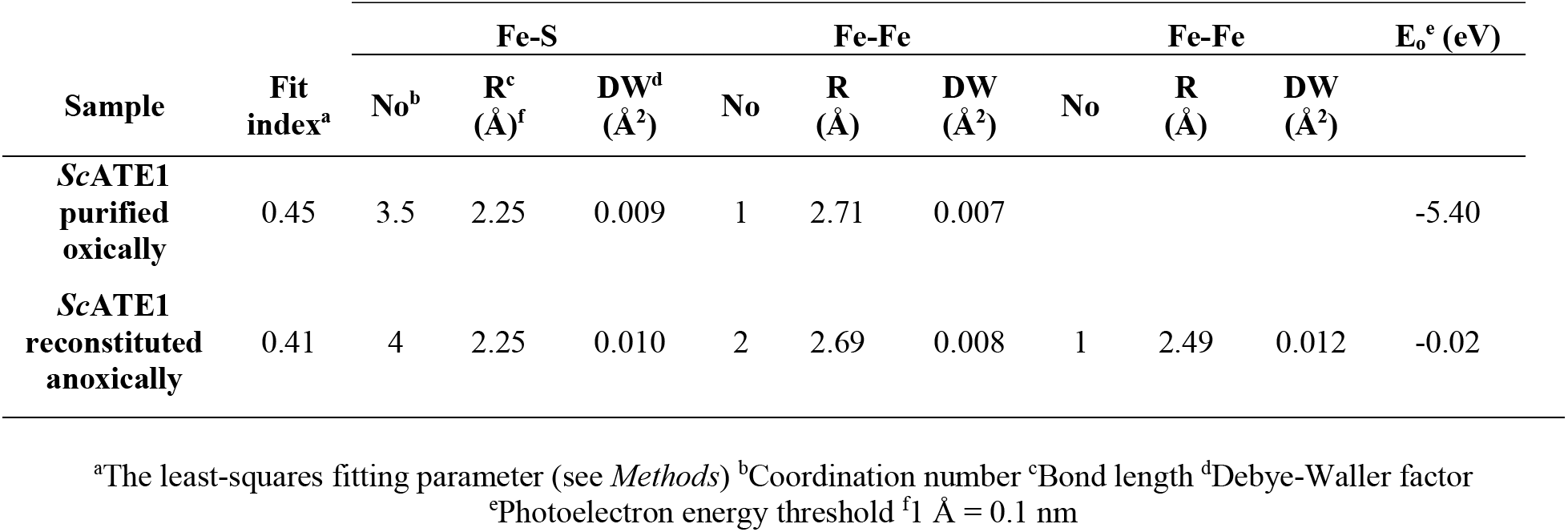
Fits obtained for the Fe K-EXAFS *Sc*ATE1 purified under oxic conditions and containing a [2Fe-2S] cluster, and *Sc*ATE1 reconstituted under anoxic conditions and containing a [4Fe-4S] cluster. Fits were determined by curve fitting using the program EXCURVE 9.2.

To further corroborate the oxidation state of the [2Fe-2S] cluster, we characterized *Sc*ATE1 expressed in the presence of ISC and isolated under oxic conditions using continuous-wave (CW) X-band electron paramagnetic spectroscopy (EPR). Consistent with our oxidation state assignment of the [2Fe-2S]^2+^ cluster based on XAS, we observe an EPR-silent spectrum under these conditions (Supplementary Figure 3a; manifested by two antiferromagnetically-coupled Fe^3+^ ions). We then added 1 mmol L^−1^ (final concentration) of either sodium ascorbate or sodium dithionite under strict anoxic conditions to the protein to reduce the cluster to the [2Fe-2S]^+^ state. In the presence of sodium ascorbate, we observed no change in the EA spectrum or the EPR spectrum. In contrast, in the presence sodium dithionite, the EA spectrum was slowly altered and effectively bleached slowly over a course of *ca*. 60 min (Supplementary Figure 3b). We then used both the ascorbate- and dithionite-reacted species in an attempt to find an EPR signal of the *S=*½ [2Fe-2S]^+^ state of the *Sc*ATE1 cluster. Despite exhaustive attempts across multiple microwave powers and multiple cryogenic temperatures (4 K to 100 K), we were unable to detect a diagnostic signal of any reduced cluster. Given the slow reaction to even the high-potential reductant sodium dithionite, and the lack of a distinct *S=*½ [2Fe-2S]^+^ EPR species, we surmise that reduction of the [2Fe-2S]^2+^ form of *Sc*ATE1 produces an unstable species that results in cluster decomposition. This hypothesis is supported by the observation that anoxic desalting of the reduced protein and subsequent exposure of *Sc*ATE1 to oxygen did not restore the diagnostic EA signal of the [2Fe-2S]^2+^ cluster.

### *Sc*ATE1 binds higher-order [Fe-S] clusters under anoxic conditions

Given that many [Fe-S] clusters can be sensitive to the presence of oxygen, and that the *Sc*ATE1 cluster appeared to be redox unstable, we then considered whether *Sc*ATE1 might be capable of binding a higher-order cluster (*e.g.*, [3Fe-4S] or [4Fe-4S]), which we tested by reconstitution under anoxic conditions. When we titrated a maximum of 10 mole eq. of Fe^3+^ and 10 mole eq. of S^2−^ over a several hour period, we noted heavy precipitation of excess iron sulfide. After gel filtration, the absorption spectrum of *Sc*ATE1 reconstituted with excess cluster components suggested the presence of a distinctly different [Fe-S] cluster than observed under oxic conditions, although there was a broad background absorption indicative of residual spuriously-bound iron sulfide (data not shown). To circumvent this problem, we titrated *Sc*ATE1 with a maximum 4 mole eq. of Fe^3+^ and 4 mole eq. of S^2−^ over a period of ≈ 2 hrs, which dramatically reduced the amount of precipitated iron sulfide and spurious binding. After filtration and buffer exchanging, determination of acid-labile iron indicated ≈ 3 eq. of Fe/polypeptide (2.68 ± 0.39; *n* = 5). The EA spectrum of *Sc*ATE1 reconstituted under anoxic conditions was most consistent with the presence of a [4Fe-4S] cluster (Fig. 3a), as only one major absorption band was present and centered at *ca*. 405 nm, common for [4Fe-4S]^2+^ clusters.^47–49^ We further confirmed this assignment using XAS and paramagnetic NMR spectroscopies. XAS again showed a first inflection point of the edge at ≈ 7120 eV and an intense pre-edge feature consistent with tetrahedral Fe centers (Supplementary Figure 4). The EXAFS data (Fig. 3b,c) were fitted with all S-based environments as the nearest neighbor ligands with an Fe-S distance of 2.25 Å (Fig. 3b,c; Table 1). In addition, the best long-range scattering interactions required multiple Fe-Fe vectors (2 longer and 1 shorter) that were fitted to distances of 2.69 Å and 2.49 Å, respectively (Fig. 3b,c; Table 1), wholly consistent with a [4Fe-4S] cubane-like cluster. The EPR spectrum of *Sc*ATE1 reconstituted under anoxic conditions was again EPR-silent, indicating the [4Fe-4S] cluster was likely in the 2+ oxidation state, as is the common oxidized state of [4Fe-4S] clusters. Finally, to confirm the cluster’s [4Fe-4S]^2+^ identity, we took advantage of paramagnetic nuclear magnetic resonance (NMR) spectroscopy, which is capable of fingerprinting the unique compositions and oxidations states of [Fe-S] clusters due to paramagnetic Cys βCH_2_ hyperfine shifts (Fig. 3d).^50,51^ The 1D ^1^H NMR spectrum of proteins containing [4Fe-4S]^2+^ clusters demonstrates 3 distinct and characteristic ^1^H shifts that are weak, located in the 10 ppm to 20 ppm region, and characteristic of this cluster composition.^50,51^ Consistent with these established characteristics, we observe three weak hyperfine shifts *ca*. 11 ppm, 15 ppm, and 18 ppm in the 1D ^1^H NMR spectrum of *Sc*ATE1 reconstituted under anoxic conditions (Fig. 3d). When considered in combination with our other spectral data, it becomes clear that *Sc*ATE1 is capable of binding a [4Fe-4S]^2+^ cluster.

**Fig. 3.**
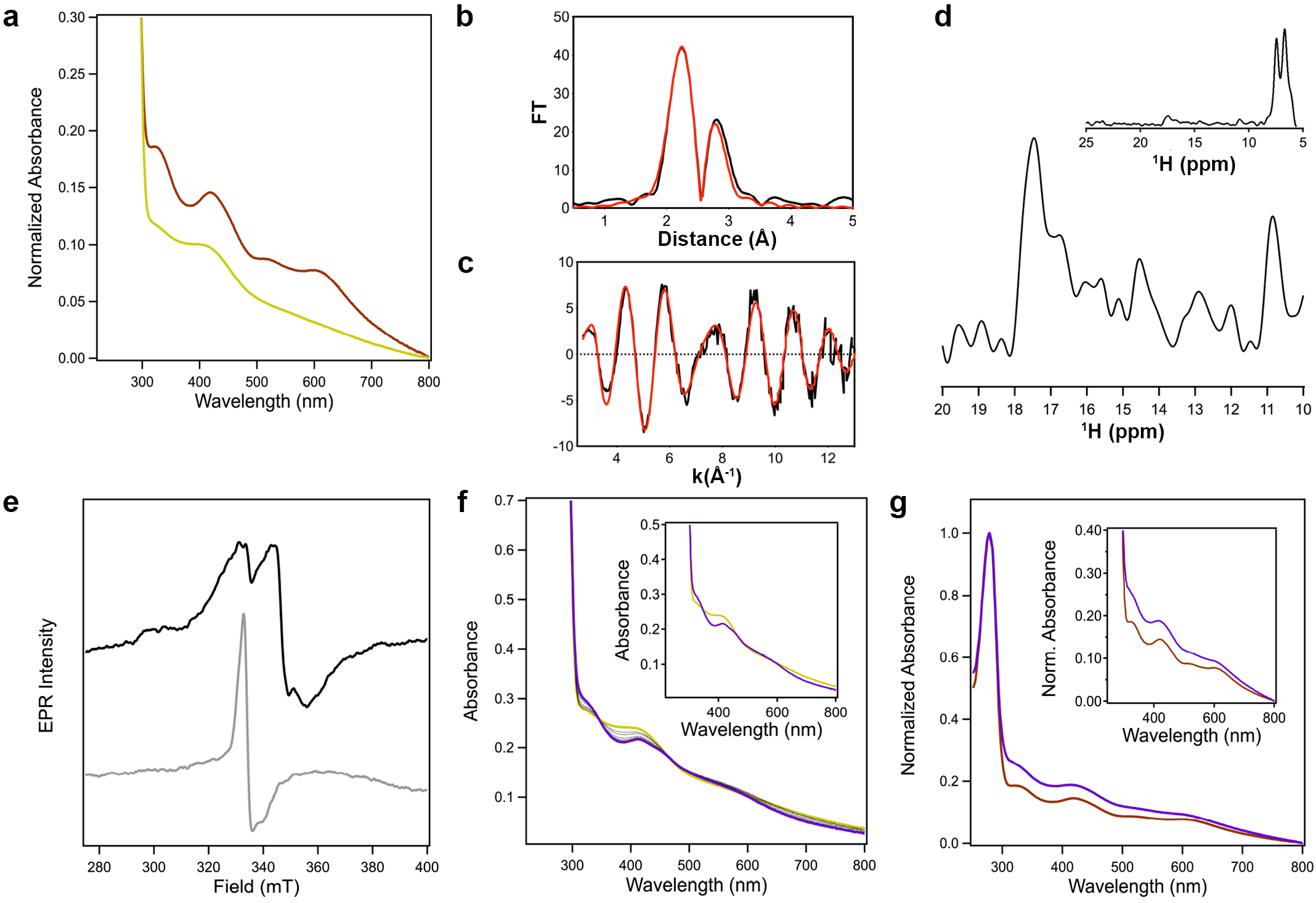
Characterization and reactivity of the [4Fe-4S] cluster of *Sc*ATE1. **a**. Reconstitution of *Sc*ATE1 under anoxic conditions yields an electronic absorption spectrum (yellow) consistent with the presence of a [4Fe-4S] cluster that is distinct from the [2Fe-2S] cluster obtained under oxic conditions (brown). **b**,**c**. EXAFS of *Sc*ATE1 reconstituted under anoxic conditions. The Fourier transformed data (**b**) and raw EXAFS (**c**) are consistent with the presence of a [4Fe-4S] cluster. 1 Å = 0.1 nm. **d**. Paramagnetic 1D 1H NMR data reveal three weak hyperfine shifts at 10.8 ppm, 14.5 ppm, and 17.5 ppm in the spectrum of *Sc*ATE1 reconstituted under anoxic conditions, consistent with a [4Fe-4S]^2+^ cluster. Inset shows the entire 1D 1H NMR region from 5-25 ppm. **e**. In the absence of sodium dithionite, an axial EPR spectrum is present that reveals the presence of some degraded [3Fe-4S]+ cluster (gray), while a [4Fe-4S]^2+^ cluster is clearly observed in the dithionite-reduced EPR spectrum (black) of anoxically purified *Sc*ATE1. Conditions were the same for both samples: frequency = 9.38 GHz, temperature = 6 K, modulation amplitude = 0.5 mT, modulation frequency = 100 kHz, 1024 points, conversion time = 175.78 ms, microwave power = 1.9 mW, 16 scans. **f**. Electronic absorption spectra reveal that the [4Fe-4S]^2+^ cluster of *Sc*ATE1 reconstituted under anoxic conditions is reactive to O_2_. The starting spectrum is yellow, the ending spectrum is purple, and the dotted black spectra represent scans taken 1 minute apart over 10 minutes. Inset: comparison of the starting (yellow) and ending (purple) spectra. **g**. Spectral comparisons reveal that the O_2_-reacted form of the [4Fe-4S]^2+^ cluster (purple) is identical to the [2Fe-2S]^2+^ cluster present in *Sc*ATE1 isolated under oxic conditions (brown).

Next, to test whether the [4Fe-4S] or the [2Fe-2S] cluster were present *in vivo*, we sought to purify *Sc*ATE1 under completely anoxic conditions after expression in anaerobic *E. coli*. To do so, we co-transformed *Sc*ATE1 pET-21a(+) into *E. coli* cells in which the chromosomal copy of transcriptional repressor protein of the *isc* operon was deleted (Δ*iscR*). Due to high constitutive levels of expressed ISC machinery, this approach is a proven method to improve *in vivo* [Fe-S] cluster insertion under anaerobic growth conditions.^43,52^ Bacteria were aerobically cultivated, then transferred to a glovebox to induce expression. Cells were subsequently harvested and lysed under an inert atmosphere (Ar_(g)_), lysed, and subjected to anoxic purification. The yield of anoxically purified *Sc*ATE1 was significantly diminished compared to oxic purification but exhibited good final purity (Supplementary Figure 5, inset). Importantly, the EA spectrum of the *Sc*ATE1 expressed and purified under anoxic conditions exhibited discrete absorption maxima at *ca*. 330 nm and *ca*. 405 nm, which closely resembles a mixture of [3Fe-4S] and [4Fe-4S] clusters (Supplementary Figure 5).^47–49,53,54^ We surmise that this mixture is the result of partial or incomplete cluster transfer to *Sc*ATE1 during anaerobic cell cultivation and heterologous protein production, or due to extreme O_2_ sensitivity in the presence of a minute amount of residual O_2_ during anoxic purification. To confirm these assignments, we collected the CW X-band EPR spectra of *Sc*ATE1 expressed and purified under anoxic conditions, both with and without the presence of sodium dithionite. In the absence of dithionite, we observed two major signals after subtraction of the background spectrum (*i.e.*, buffer only): one sharp signal at *g* ≈ 4.30 consistent with adventitious or misloaded Fe^3+^, and a second axial signal with *g*_∥_ ≈ 2.00 and *g*_⊥_ ≈ 1.97 consistent with a [3Fe-4S]^+^ *S*=½ species (Fig. 3e).^54,55^ In the presence of dithionite and after subtraction of the background spectrum (*i.e.*, buffer with dithionite), we observed only a rhombic signal with *g*_1_ ≈ 2.03, *g*_2_ ≈ 1.93, and *g*_3_ ≈ 1.88, consistent with a [4Fe-4S]^+^ *S*=½ species (Fig. 3e).^47,56,57^ Thus, when our anoxic reconstitution and purification results are taken together, it becomes evident that *Sc*ATE1 binds a higher-order cluster in the absence of oxygen and *in vivo*.

### The *Sc*ATE1 [Fe-S] cluster is oxygen-sensitive

Given the differences in the [Fe-S] cluster identity as a function of oxic *vs*. anoxic purification of *Sc*ATE1, we hypothesized that the ATE1 cluster might be oxygen-sensitive. We confirmed this hypothesis two different ways. First, after anoxic reconstitution and buffer exchanging to remove excess iron and sulfide, exposure of the reconstituted [4Fe-4S]^2+^ to ambient atmosphere resulted in rapid (*t*_1/2_ ≪ 5 min) and isosbestic conversion to a different species (Fig. 3f). Second, we performed the same experiment with *Sc*ATE1 that was produced and purified under anoxic conditions (mixture of [3Fe-4S]^+^ and [4Fe-4S]^2+^ clusters) and noted nearly identical behavior on a very similar timescale (Supplementary Figure 6). In both cases, the final EA spectrum exhibited multiple peaks that were virtually superimposable with that of EA spectrum of *Sc*ATE1 isolated under oxic conditions (Fig. 3g), pointing to the presence of a [2Fe-2S]^2+^ cluster as a common species generated upon exposure of the higher-order cluster to O_2_. These results demonstrate that the *Sc*ATE1 [Fe-S] cluster is oxygen-sensitive, which could be leveraged to sense oxidative stress within the cell in order to alter arginylation efficacy and/or substrate specificity.

### The *Sc*ATE1 [Fe-S] cluster is bound in an evolutionarily-distinct N-terminal domain

To determine the binding location of the [Fe-S] cluster, we first used bioinformatics data on known ATE1 protein sequences to pinpoint Cys residues involved in cluster binding. Partial sequence alignments of ATE1s from a continuum of organisms (yeast to human) reveal strong conservation of 4 Cys residues (numbered based on *Sc*ATE1): Cys^20^/Cys^23^ (forming a CxxC motif), and Cys^94^/Cys^95^ (forming a CC motif) (Supplementary Figure 7). These Cys residues are clustered within the N-terminal region of the ATE1 polypeptide, which has an unknown function and is distinct from bacterial leucyl/phenylalanyl-(L/F)-tRNA protein transferases.^4^ Additional sequence context indicates that this region is the location for cluster binding. Recent work has shown that an LYR-motif (variations including “LYK” and “VYK”) is an important polypeptide sequence indicator of eukaryotic [Fe-S] cluster locations. This key amino acid signature engages the co-chaperone HSC20, guiding the binding and the delivery of [Fe-S] cluster maturation machinery in several eukaryotic organisms.^58–61^ Importantly, sequence alignments demonstrate the strong conservation of a LYR/LYK/VYK-like motif in sequence proximity along the ATE1 N-terminal extension (Supplementary Figure 7). Combined, these data indicated that the ATE1 N-terminal domain could have a function in [Fe-S] cluster binding.

To test this hypothesis, we first cloned the N-terminal domain of *Sc*ATE1 (residues 1-146) into a pET vector and attempted to express this domain in *E. coli*. Despite exhaustive efforts and expression testing using multiple cell lines and multiple media types, the tagged N-terminal domain failed to accumulate, suggesting this domain may be unstable on its own. Instead, we generated a construct of N-terminal *Sc*ATE1 fused to maltose-binding protein (MBP *Sc*ATE1-N; Supplementary Figure 8a), an effective strategy in previous methods for recombinant production of small proteins/domains that bind [Fe-S] clusters.^47^ We then purified this fusion protein (Supplementary Figure 8b) and subjected it to chemical reconstitution under anoxic conditions. After anoxic desalting, the MBP *Sc*ATE1-N retained a similar Fe/polypeptide stoichiometry, and the electronic absorption spectrum was nearly superimposable with that of *Sc*ATE1 reconstituted under anoxic conditions (Supplementary Figure 8c). These results confirm that the N-terminal domain is the site of cluster binding in *Sc*ATE1, which is wholly consistent with very early site-directed mutagenesis studies on *Sc*ATE1 that demonstrated that arginylation efficacy was linked to four strongly-conserved N-terminal Cys residues (Cys^20/23/94/95^) (*vide infra*).^62^

### The *Sc*ATE1 [Fe-S] cluster affects *in vitro* arginylation

To determine whether the presence of the [Fe-S] cluster affected arginylation efficacy, we first tested the activity of our apo *Sc*ATE1 expressed in the absence of cluster building blocks (*i.e.*, ISC, exogenous ferric citrate, and exogenous *L*-Cys). We used a mass spectrometry-based assay that takes advantage of the natural Glu residue on the N-terminus of α-lactalbumin (ALB),^63^ allowing us to do a direct, top-down mass spectrometric comparison between the non-arginylated (molecular mass = 14176 g/mol) and the arginylated ALB (molecular mass = 14333 g/mol). *Sc*ATE1 produced in the absence of ISC and without media supplementation (≤ 0.1 Fe ion per polypeptide) was active towards arginylation in the presence of recombinant *E. coli* arginine synthetase (ArgRS) and both *E. coli* and yeast tRNA^Arg^ (Fig. 4a,b). This reaction was evidenced by the increase in m/Z ratio of observed ALB after reaction, which corresponds to a mass difference of 156 g/mol (Fig. 4b): plus one Arg residue (174 g/mol) minus one H_2_O (18 g/mol) upon peptide bond formation after calculation of the peptide charge (Z=14). This N-terminal modification is dependent on the presence of *Sc*ATE1, as no arginylation is observed in the absence of *Sc*ATE1 (Fig. 4b,c). These results indicate that our recombinant *Sc*ATE1 is enzymatically competent in the apo form. We then tested for arginylation of ALB in the presence of *Sc*ATE1 with an [Fe-S] cluster. Because of the rapid oxygen-sensitive nature of the [3Fe-4S]^+^- and [4Fe-4S]^2+^-bound forms of *Sc*ATE1, we could not assess the arginylation efficacy and confidently guarantee cluster composition. However, we were able to assess the effects of the oxygen-stable [2Fe-2S]^2+^-bound *Sc*ATE1 on ALB arginylation. Importantly, we observed a *ca*. 2-fold increase in the ability of the [2Fe-2S]^2+^-bound *Sc*ATE1 to arginylate ALB when compared to apo *Sc*ATE1 under identical conditions (Fig. 4d; average of 39.7 % arginylation of [2Fe-2S]^2+^-bound *Sc*ATE1 compared to an average of 20.6 % arginylation of apo *Sc*ATE1). Although, there are numerous potential substrates that remain to be screened, these exciting results indicate that the presence of the O_2_-reacted [Fe-S] cluster functions as a positive regulator of arginylation.

**Fig. 4.**
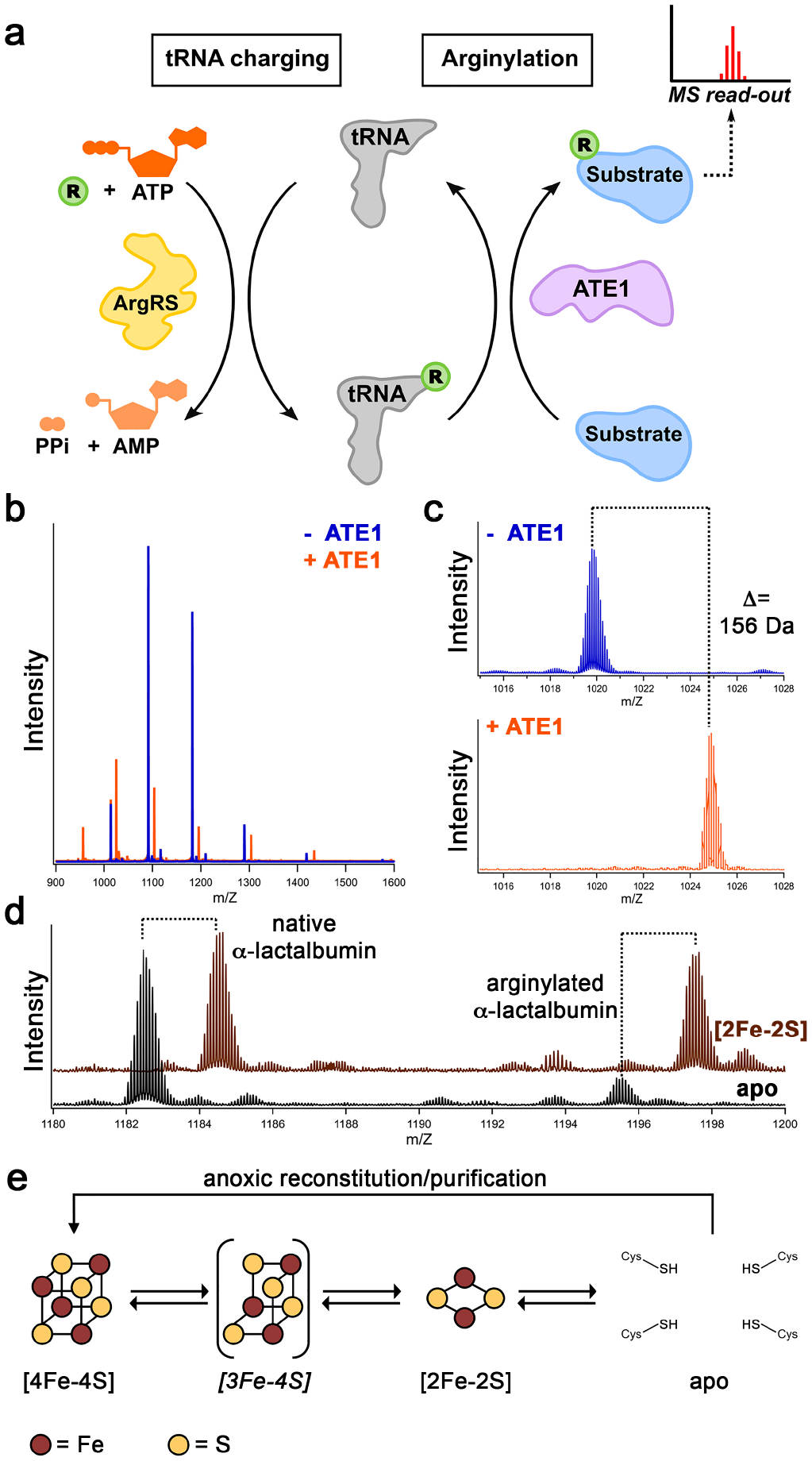
*Sc*ATE1-catalyzed arginylation is affected by the [Fe-S] cluster. **a**. Cartoon schematic of the recombinant arginylation assay. The arginylated tRNA (Arg tRNA^Arg^) is catalyzed by arginine synthetase (ArgRS) aminoacylation. Arginylation of substrate is catalyzed by ATE1, which is characterized by mass spectrometry. **b**. The arginylation of a model substrate (α-lactalbumin) is *Sc*ATE1-dependent, as evidenced by the shift in the top-down mass spectral envelope of α-lactalbumin in the absence of *Sc*ATE1 (blue) and the presence of *Sc*ATE1 (orange). **c**. Close-up of the mass spectral window from 1014 m/Z to 1028 m/Z. The change in molecular weight (Δ=156 g/mol (Da)) is consistent with a single arginylation event after considering the charge (+14) of the polypeptide. **d**. In the presence of the [2Fe-2S]^2+^ cluster-bound form of *Sc*ATE1 (brown spectrum), the extent of arginylation of α-lactalbumin is increased *ca*. 2-fold on average compared to apo *Sc*ATE1 (black spectrum). **e**. Cartoon synopsis of the cluster-bound forms of *Sc*ATE1. The [4Fe-4S]- and [2Fe-2S]-cluster bound forms may be interconverted in an O_2_-dependent manner, presumably through a [3Fe-4S] intermediate (bracketed, as it has not yet been observed under these conditions). We believe a [4Fe-4S] cluster is operative under anoxic conditions in the cell until reaction with O_2_ converts the cluster to [2Fe-2S], and we observe such behavior when *Sc*ATE1 is purified and/or reconstituted under anoxic conditions and subsequently exposed to O_2_.

### Heme binding induces ATE1 aggregation and inactivation

We next sought to rectify our observations that *Sc*ATE1 binds an [Fe-S] cluster with the previous implication that ATE1 could be a heme-binding protein. Despite our efforts using media supplementation, under no circumstances did we observe incorporation of heme into *Sc*ATE1 (*vide supra*). We then attempted *in vitro* reconstitution by titrating hemin into *Sc*ATE1. Over the course of adding 1 mole eq. to 5 mole eq. hemin, we commonly observed visible protein precipitation. Once centrifuged, this protein pellet was colored, indicating an unknown amount of co-precipitation of hemin and *Sc*ATE1. For the fraction of protein that did not precipitate, we desalted to remove any excess hemin in solution. Intriguingly, the absorbance spectrum of the remain heme-bound *Sc*ATE1 (Supplemental Figure 9) is strikingly similar to the spectra of non-specific heme-protein interactions, as previously determined.^64^ Furthermore, we found that the pools of protein with the highest ratio of heme to *Sc*ATE1 (≈ 1, estimated by the ratio of A_280_/A_400_) appear to be aggregated, as determined by size-exclusion chromatography (SEC) (Supplemental Figure 10 a,b). In fact, this aggregation appears to be heme-dependent, and we observe a clear shift to higher-order aggregation based on SEC as we titrated hemin into *Sc*ATE1 (Supplemental Figure 10 a,b). Moreover, these pools of aggregated *Sc*ATE1 were inactive as determined by our MS assay (data not shown), and as previously observed for mouse (*Mus musculus*) ATE1^33^. Thus, these data indicate that heme binding to *Sc*ATE1 may be an off-target effect that is distinct from that of cluster-binding and results in deactivation via protein aggregation. We believe this same heme-induced aggregation may have been operative in previous studies of heme binding inactivation of *Mm*ATE1,^33^ which we believe is also an [Fe-S] protein (*vide infra*).

### The ATE1 [Fe-S] cluster is evolutionarily conserved

Finally, because our partial sequence alignments reveal strong conservation of cluster-binding motifs within the N-terminal region of ATE1s across evolutionary space (Supplementary Figure 7), we believe these observations strongly indicate [Fe-S] clusters are a common feature of eukaryotic ATE1s. To strengthen this argument, we expressed and purified N-terminally tagged *Mus musculus* ATE1 isoform 1B7A (*Mm*ATE1^1B7A^), which was previously studied in the context of heme binding^33^, and we tested this construct for cluster incorporation. Indeed, while expression of *Mm*ATE1^1B7A^ in *E. coli* Rosetta II cells grown in LB media supplemented with heme precursors and purified under oxic conditions resulted in no sign of heme incorporation, concentrated purified *Mm*ATE1^1B7A^ was also slightly brown in color (data not shown). Because of overlapping antibiotic resistance between the Rosetta II cells and the ISC machinery (both Cam^r^), we were unable to express this construct in the identical manner of *Sc*ATE1. However, we expressed *Mm*ATE1^1B7A^ in the presence of exogenous ferric citrate and *L*-Cys in the growth media, and the concentrated, purified protein exhibited an intense brown color, indicative of [Fe-S] clusters. It is clear from the EA spectrum of *Mm*ATE1^1B7A^ purified under oxic conditions that a [2Fe-2S]^2+^ cluster nearly identical to that observed in *Sc*ATE1 is present (Supplementary Figure 11), and reconstitution under anoxic conditions results in a spectrum that looks nearly identical to that of the [4Fe-4S]^2+^ cluster bound to *Sc*ATE1 and MBP *Sc*ATE1-N reconstituted under identical conditions (Supplementary Figure 11). Further characterization of the cluster-bound forms of *Mm*ATE1^1B7A^ are currently underway, but these results clearly point to [Fe-S] cluster binding as a common property of both yeast and mammalian ATE1s.

## DISCUSSION

ATE1s are critical eukaryotic enzymes that arginylate a diverse set of protein targets, which are essential for general cellular homeostasis and function of higher-order eukaryotes.^4^ The process of arginylation may regulate protein function in one of two major ways. In a non-degradative manner, arginylation may stabilize proteins or even change their activity and/or structure. For example, α-synuclein and β-amyloid, both of which are proteins implicated in neurological diseases, are stabilized and have lower aggregation propensities when arginylated.^65,66^ In another example, arginylation can change oligomerization of essential structural proteins, such as the cytoskeletal protein β-actin.^67,68^ In contrast, arginylation can also target proteins for degradation via the N-degron pathway. Once arginylated, E3 ubiquitin ligases of the ubiquitin proteasome system (UPS) ubiquitinate target polypeptides for cellular clearance. While the exact extent of the true intracellular N-degron arginylome remains to be uncovered, proteins linked to the Arg N-degron pathway are implicated in the yeast stress response,^69^ plant growth,^26^ and even mammalian cardiovascular^9,22^ and neurological^66^ development. Thus, the fidelity of post-translational arginylation, which may be achieved through multiple levels and types of regulatory processes, is paramount.

Previous results have indicated that ATE1-connected pathways may be subject to different methods of regulation such as small diatomic molecules, cofactors, and even protein-protein interactions. In plants, it is now well-established that the response to hypoxia/O_2_ is connected to the N-degron pathway.^34,70,71^ Group VII ethylene response factors (ERFs)—proteins that regulate the plant hypoxic response—were shown to be targets of the N-degron pathway in an O_2_-dependent manner. Work has demonstrated that several of the group VII ERFs have N-terminal Met-Cys sequences, and ERF protein degradation is linked to the oxidation of the N-terminal Cys residue^72^ (presumably after aminopeptidase cleavage) dependent on oxygen availability and the enzymatic capabilities of plant CDOs.^20^ These findings connect the concentration of intracellular O_2_ and the N-degron pathway in plants. A similar connection has been made between NO availability and the N-degron pathway in mammals, where NO has been shown to oxidize N-terminal Cys residues directly rather than enzymatically, rendering secondary destabilizing residues that are recognized by ATE1 for arginylation.^21^ NO-linked oxidation of N-terminal Cys residues has been shown to have dramatic effects on the degradation of regulators of G-protein signaling (RGSs), which are critical for cardiovascular maturation in mammals.^21^ Although these are indirect regulatory responses (occurring at the protein substrate that is the target of arginylation), one study has shown that *Mm*ATE1^1B7A^ may be inactivated directly by the presence of ferric heme through the formation of a strained disulfide, which presumably alters protein structure.^33^ Furthermore, ATE1 may be upregulated through direct protein-protein interactions with a partner protein present in higher-order eukaryotes known as Liat1.^73^ The molecular details of this protein-protein interaction remain to be fully described; however, it was demonstrated that binding of this small protein to certain ATE1 isoforms increased the arginylation of a model ATE1 substrate *in vitro*.^73^ Thus, given the multi-layered control of the Arg N-degron pathway, it is perhaps unsurprising that ATE1 itself may also bind and utilize a regulatory cofactor.

In this work, we have discovered that ATE1s bind [Fe-S] clusters, expanding the regulatory complexity of the Arg N-degron pathway. Our initial focus on the heme-binding properties of ATE1 revealed instead that ATE1 binds an [Fe-S] cluster. Purified under oxic conditions, our data revealed the presence of a [2Fe-2S]^2+^ cluster (Fig. 4e), which has increased activity towards the *in vitro* N-terminal arginylation of a model ATE1 substrate. To determine if a higher-order cluster could form, anoxic reconstitution of ATE1 yielded a characteristic [4Fe-4S]^2+^ cluster (Fig. 4e), and these results were recapitulated by anoxic purification of ATE1, indicating that a [4Fe-4S]^2+^ cluster could be formed *in vivo* (in addition to a partially-loaded [3Fe-4S]^+^ cluster; Fig. 4e). In either case, the cluster is oxygen-sensitive, displaying rapid decomposition from [4Fe-4S]^2+^ to [2Fe-2S]^2+^ (presumably through a [3Fe-4S]^+^ intermediate; Fig. 4e) upon exposure to O_2_. We subsequently mapped the location of the oxygen-sensitive ATE1 [Fe-S] cluster to the N-terminal domain, which is evolutionarily-conserved, contains four key Cys residues that are important for arginylation activity, and contains an LYK-like motif that is known to govern [Fe-S]-cluster loading in eukaryotes.^58–61^ Consistent with our hypothesis that cluster-binding is a common feature of ATE1s, we demonstrated that *Mm*ATE1^1B7A^ is also capable of binding an [Fe-S] cluster. Given these results, we propose that an ATE1 [Fe-S] cluster is present in all eukaryotes, including humans.

It is tempting to speculate that the presence of the ATE1 [Fe-S] cluster and its O_2_-sensitivity may be linked to function, given the capabilities of the N-degron pathway to sense NO and O_2_ (*vide supra*). The binding of an [Fe-S] cluster by ATE1 would be well matched to these known functions of the N-degron pathway, as [Fe-S] proteins are either known or speculated to sense O_2_, NO, and even the redox state of the cell.^47,48,74–76^ We do not have evidence that this cluster functions as an electron transport site and there is no need for electron transport in the arginylation reaction (Fig. 4a); furthermore, reduction of the cluster in ATE1 is relatively slow and results in instability and loss of cluster loading. Instead, we posit that the evolutionarily-conserved [Fe-S] cluster within the ATE1 N-terminal domain may function to sense O_2_, NO, and even respond to oxidative stress. Under such conditions, N-terminal Cys residues may be oxidized, and/or proteins may be misfolded resulting in a greater need to flag and to degrade dysfunctional polypeptides. We imagine that altered compositions of the cluster may transmit this information through the N-terminal domain to the catalytic C-terminal domain that is predicted to contain the GNAT fold, potentially changing the rate of arginylation and even arginylation specificity. This function, which warrants future in-depth studies, would be entirely consistent with previous results demonstrating that the Arg N-degron pathway is capable of sensing and responding to changes in intracellular O_2_ and NO levels.

We believe that our proposed [Fe-S] cluster-based sensor hypothesis is distinct from previous observations of hemin interactions with ATE1. The prior studies of *Mm*ATE1^1B7A^ mapped the ATE1-heme interaction to residue Cys^411^,^33^ located within the C-terminal domain of ATE1 and distinct from the Cys residues involved in [Fe-S] coordination. However, it is not uncommon for exposed Cys residues to interact with heme in an adventitious manner,^64^ and *Mm*ATE1^1B7A^ appeared to harbor multiple potential heme-interacting sites with very modest binding affinities.^33^ The stated causal relationship of decreased arginylation of *Mm*ATE1^1B7A^ was hemin-derived oxidation of two adjacent Cys residues to an unusually strained disulfide in the N-terminal domain, but it is difficult to rectify how this occurs via heme binding in the C-terminal catalytic region. It is commonly accepted that there is no free heme within the cell under homeostatic conditions (*i.e.*, heme is either bound tightly by hemoproteins, heme chaperones, or heme scavengers),^77^ but it is possible that heme may be spuriously released under dyshomeostasis during oxidative stress or infection within the cell, a situation that could lead to the observed decrease in ATE1 activity. Contrary to previous work, we find that ATE1 appears to bind heme non-specifically and induces protein aggregation leading to inactivation, suggesting the heme-based inactivation may be irrelevant to ATE1’s native function. On the basis of our results, we present the compelling discovery that [Fe-S] clusters are the operative cofactors that bind and give function to the intriguing ATE1 N-terminal domain. This discovery opens up many new avenues of exploration within this family of essential eukaryotic enzymes.

## Acknowledgements

This work was supported by NIH-NIGMS grant R35 GM133497 (A.T.S) and NIH-NIGMS grant T34 GM136497 (N.-E.E.). In addition, this work was financially supported by the Deutsche Forschungsgemeinschaft (DFG) Priority Programme “Iron–Sulfur for Life: Cooperative Function of Iron–Sulfur Centers in Assembly, Biosynthesis, Catalysis and Disease” (SPP 1927) IS 1476/4-1 (I.S.); the Chemical Industry Fund Li 196/05 (I.S.) and Hoe 700080 (H.R.); and the German Academic Scholarship Foundation (H.R.). Sequence searches utilized both database and analysis functions of the Universal Protein Resource (UniProt) Knowledgebase and Reference Clusters (http://www.uniprot.org) and the National Center for Biotechnology Information (http://www.ncbi.nlm.nih.gov/). Use of the Stanford Synchrotron Radiation Lightsource, SLAC National Accelerator Laboratory, is supported by the U.S. Department of Energy, Office of Science, Office of Basic Energy Sciences under Contract No. DE-AC02-76SF00515. The SSRL Structural Molecular Biology Program is supported by the DOE Office of Biological and Environmental Research, and by the National Institutes of Health, National Institute of General Medical Sciences (including P41 GM103393).

## Author contributions

V.V., J.B.B., H.R., I.M., K.N.C., V.S., and A.T.S. designed the research; V.V., J.B.B, H.R., I.M.,N.-E.E., T.S.B., V.S., K.N.C., and A.T.S. performed the research; V.V., J.B.B., H.R., V.S., K.N.C., I.S., and A.T.S. analyzed the data; and V.V., J.B.B., H.R., I.M., V.S., K.N.C., I.S., and A.T.S. wrote and edited the paper.

## Additional information

### Supplementary information

Supplementary information accompanies this paper.

### Competing interests

The authors declare no competing interests.

## METHODS

### Materials

Modified forms of the BL21(DE3) and C43(DE3) *E. coli* expression cell lines in which the gene for the multidrug exporter AcrB (a common *E. coli* contaminant) had been deleted (BL21(DE3) Δ*acrB* and C43(DE3) Δ*acrB*, respectively) were generous gifts of Prof. Edward Yu (Case Western Reserve University). All materials used for buffer preparation, protein expression, and protein purification were purchased from commercial vendors and were used as received.

### Cloning, expression, and aerobic purification of *Sc*ATE1 constructs

DNA encoding for the gene corresponding to the single-isoform arginine transferase (ATE1) from *Saccharomyces cerevisiae* (strain ATCC 204508; Uniprot identifier P16639) (*Sc*ATE1) was commercially synthesized with an additionally engineered DNA sequence encoding a C-terminal TEV-protease cleavage site (ENLYFQS). This gene was subcloned into the pET-21a(+) expression plasmid using the NdeI and XhoI restriction sites, encoding a C-terminal (His)_6_ affinity tag when read in-frame. The complete expression plasmid was transformed into chemically competent BL21(DE3) Δcells, spread onto Luria-Bertani (LB) agar plates supplemented with 100 μg/mL ampicillin, and grown overnight at 37 °C. Colonies from these plates served as the source of *E. coli* for small-scale starter cultures (generally 100 mL LB supplemented with 100 μg/mL ampicillin). Large-scale expression of *Sc*ATE1 was accomplished in 12 baffled flasks each containing 1 L sterile LB supplemented with 100 μg/mL (final) ampicillin and inoculated with a pre-culture. Cells were grown by incubating these flasks at 37 °C with shaking until the optical density at 600 nm (OD_600_) reached ≈ 0.6 to 0.8. The flasks containing cells and media were then chilled to 4 °C for 2 h, after which protein expression was induced by the addition of isopropyl β-D-l-thiogalactopyranoside (IPTG) to a final concentration of 1 mmol/L. The temperature of the incubator shaker was lowered to 18 °C with continued shaking. After *ca*. 18 h to 20 h, cells were harvested by centrifugation at 4800×*g*, 10 min, 4 °C. Cell pellets were subsequently resuspended in resuspension buffer (50 mmol/L Tris, pH 7.5, 100 mmol/L NaCl, 0.7 mol/L glycerol), flash-frozen on N_2(l)_, and stored at −80 °C until further use.

A similar approach was taken to generate constructs of the *Sc*ATE1-N terminal domain (*Sc*ATE1-N) and a maltose-binding protein fusion of the *Sc*ATE1-N terminal domain (MBP *Sc*ATE1-N). DNA encoding for the genes of the *Sc*ATE1 N-terminal domain (residues 1-146), with additionally engineered DNA sequences encoding for a C-terminal TEV-protease cleavage site (ENLYFQG) or with an additionally engineered DNA sequence encoding for an N-terminal maltose-binding protein sequence (based on P0AEX9: *Escherichia coli* (K-12) *malE* gene product) followed by a Tobacco Etch Virus (TEV)-protease cleavage site, was synthesized. For the former approach, the gene was subcloned into the pET-21a(+) expression plasmid using the NdeI and XhoI restriction sites, encoding for a C-terminal (His)_6_ affinity tag when read in-frame. For the latter approach, gene was subcloned into the pET-45b(+) expression plasmid using the PmlI and PacI restriction sites, encoding for a N-terminal (His)_6_ affinity tag followed by maltose-binding protein when read in-frame. Transformations, cell growths, and cell resuspensions for both constructs were identical to those of the WT construct.

All steps for the purification of *Sc*ATE1 were performed at 4 °C unless otherwise noted. Briefly, frozen cells were thawed and stirred until the solution was homogeneous. Solid phenylmethylsulfonyl fluoride (PMSF; 50 mg to 100 mg), and a solution of tris(2-carboxyethyl)phosphine hydrochloride (TCEP; 1 mmol/L final concentration) were added immediately prior to cellular disruption using an ultrasonic cell disruptor set to a maximal amplitude of 70 %, 30 s pulse on, 30 s pulse off, for a total pulse on time of 12 min. Cellular debris was cleared by ultracentrifugation at 163000×*g* for 1 h. The supernatant was then applied to a 5 mL metal affinity column that had been charged with Ni^2+^ and equilibrated with 8 column volumes of wash buffer (50 mmol/L Tris, pH 8.0, 300 mmol/L NaCl, 1 mmol/L TCEP, and 1.4 mol/L glycerol) with an additional 21 mmol/L imidazole. The column was then washed with 12 column volumes (CVs) of wash buffer with an additional 30 mmol/L imidazole. Protein was then eluted with wash buffer containing an additional 300 mmol/L imidazole. Fractions were concentrated using a 15 mL 30 kg/mol (30 kDa) molecular-weight cutoff (MWCO) spin concentrator. Protein was then applied to a 120 mL gel filtration column that had been pre-equilibrated with 50 mmol/L Tris, pH 7.5, 100 mmol/L KCl, 1 mmol/L dithiothreitol (DTT), and 0.7 mol/L glycerol. The eluted fractions of the protein corresponding to the protein monomer (≈ 60000 g/mol; ≈ 60 kDa), were pooled and concentrated with a 4 mL 30 kg/mol (30 kDa) MWCO spin concentrator. Protein concentration was determined using the Lowry assay, and purity was assessed via SDS-PAGE analysis (acrylamide mass fraction of 15 %).

To purify MBP *Sc*ATE1-N, a similar protocol was followed for the WT construct with the following modifications. The supernatant was applied to two tandem 5 mL amylose columns that had been pre-equilibrated with 5 CVs of wash buffer (25 mmol/L Tris, pH 7.5, 200 mmol/L NaCl, 0.7 mol/L glycerol, 1 mmol/L TCEP). The column was then washed with 20 CVs of wash buffer. Protein was then eluted by wash buffer containing 10 mmol/L maltose. Fractions were concentrated using a 15 mL 30 kg/mol (30 kDa) MWCO spin concentrator. Protein concentration was determined using the Lowry assay, and purity was assessed via SDS-PAGE (acrylamide mass fraction of 15 %) analysis.

To express *Sc*ATE1 in the presence of heme and/or iron-sulfur precursors, the protein was expressed as previously described (*vide supra*) with some modifications. To encourage the incorporation of heme, immediately prior to addition of IPTG, δ-aminolevulinic acid (1 mmol/L final concentration) and ferric citrate (125 μmol/L final concentration) were added to the expression cultures. To encourage the presence of iron-sulfur clusters, immediately prior to addition of IPTG, *L*-Cys (167 mg/L final concentration) and ferric citrate (125 μmol/L final concentration) were added to the expression cultures. All subsequent steps were performed as previously described (*vide supra*).

To express *Sc*ATE1 in the presence of the *E. coli* iron-sulfur cluster (ISC) biosynthesis machinery, the *Sc*ATE1 expression plasmid (encoding for ampicillin resistance) was transformed into chemically-competent C43 (DE3) Δ*acrB* cells bearing the pACYC184iscS-fdx plasmid (encoding for chloramphenicol resistance and constitutively expressed), spread onto LB agar plates supplemented with 100 μg/mL ampicillin and 34 μg/mL chloramphenicol, and grown overnight at 37 °C. Large-scale expression was accomplished as above with the following modifications: immediately prior to addition of IPTG, *L*-Cys (167 mg/L final concentration) and ferric citrate (125 μmol/L final concentration) were added to the expression cultures. After *ca*. 18 h to 20 h, cells were harvested, lysed, and protein was purified as previously described. There was no dramatic change in protein yield, purity, or homogeneity; however, the purified protein was deeply colored even after size-exclusion chromatography.

Western blotting was used to confirm the presence of the (His)_6_-tagged *Sc*ATE1 constructs. Briefly, protein samples were run on SDS-PAGE (acrylamide mass fraction of 15 %) and then transferred to a PVDF membrane overnight. Blocking of the membrane was done at room temperature by addition of instant milk solution (milk mass fraction of 5 %) in phosphate-buffered saline with Tween 20 (PBST; Tween 20 volume fraction of 1 %) with rocking for 1 h. The membrane was then washed with PBST, and an anti-(His)_6_ antibody that was diluted *ca*. 1:6000 (volume fraction) into an instant milk solution (milk mass fraction of 1 %) in PBST and incubated with rocking for 1 h. The membrane was then washed with PBST and developed using a chromogenic detection of horseradish peroxidase kit.

### Anoxic expression and purification of *Sc*ATE1

Overnight starting cultures of *E. coli* BL21(DE3) *ΔiscR* containing the pET21(+) *Sc*ATE1 plasmid were used to inoculate Terrific Broth (TB) medium. TB medium was supplemented with kanamycin (50 μg/mL), ampicillin (100 μg/mL) and ferric ammonium citrate (2 mmol/L). Cells were cultivated under oxic conditions at 37 °C with shaking until OD_600_ reached ≈ 2. For anoxic cell growth cultures were then moved to an anoxic glove box containing an H_2_/N_2_ atmosphere and operating at < 7 mg/m^3^ (5 ppm) O_2_. Gene expression was induced by adding IPTG to a final concentration of 0.5 mmol/L. To facilitate [Fe-S] cluster assembly and anaerobic metabolism, 2 mmol/L *L*-cysteine as well as 25 mmol/L sodium fumarate were added. Cultures were stirred on a magnetic stirrer at RT for 20 h following induction.

Cells were harvested by spinning for 10 min at 6000×*g* and 4 °C, while cultures were covered with Ar_(g)_ to maintain anaerobic conditions. For cell lysis, cells were resuspended in 5 mL buffer A (50 mmol/L Tris, pH 8.0, 300 mmol/L NaCl, 1.4 mol/L glycerol) per gram of cell pellet. Protease inhibitor tablets were added as needed. After stirring at RT for 20 min under anaerobic conditions the Ar_(g)_ covered suspension was sonicated for 20 min with an amplitude of 60 % and a pulse of 1 s every 3 s using an ultrasonic cell disrupter. Argon-covered lysates were clarified by centrifugation at 40000×*g*.

*Sc*ATE1 purification was carried out under anoxic conditions at RT. Anoxic lysate was applied to a metal affinity column with a bed volume of 5 mL equilibrated with buffer A. The column was then washed with 10 CVs buffer A and 10 CVs of a mixture of buffers A and B in (buffer B volume fraction of 10 %, 50 mmol/L Tris, pH 8.0, 300 mmol/L NaCl, 1.4 mol/L glycerol, 300 mmol/L imidazole) before the target protein was eluted with only buffer B. Fractions containing the brown-colored target protein were pooled and subsequently applied to a 24 mL size-exclusion column pre-equilibrated in 25 mmol/L Tris, pH 7.5, 0.7 mol/L, 100 mmol/L KCl. Eluted monomeric protein was then further used for electronic absorption and electron paramagnetic spectroscopies (*vide infra*).

### Purification of *Mm*ATE1^1B7A^

An ampicillin-resistant expression plasmid encoding for mouse (*Mus musculus*) ATE1 isoform 1B7A N-terminally tagged with a (His)_10_ tag tethered to Ub (His-10-Ub-ATE1^1B7A^) was a generous gift of Prof. Alexander Varshavsky (CalTech) and Prof. Rong-Gui Hu (Shanghai Institutes for Biological Sciences).^33^ The expression plasmid was transformed into chemically competent Rosetta 2 (DE3) cells, spread onto LB agar plates supplemented with 100 μg/mL ampicillin and 34 μg/mL chloramphenicol, and grown overnight at 37 °C. Expression of *Mm*ATE1^1B7A^ was identical to that of *Sc*ATE1 (*vide supra*) except that large-scale expression flasks were supplemented with 34 μg/mL chloramphenicol in addition to 100 μg/mL ampicillin. Because the ISC plasmid bore the same antibiotic resistance as that of the Rosetta II expression cells, this commonality precluded the testing of expression of *Mm*ATE1^1B7A^ in the presence of the ISC machinery. Instead, cells were grown in the presence of supplemented *L*-Cys and ferric citrate, as described (*vide supra*). The purification of *Mm*ATE1^1B7A^ was identical to that of *Sc*ATE1 (*vide supra*).

### Cloning, Expression, and Purification of *Ec*ArgRS

DNA encoding for the gene corresponding to the Arg tRNA synthetase from *E. coli* (strain K12; Uniprot identifier P11875) (*Ec*ArgRS) was commercially synthesized with an additionally engineered DNA sequence encoding for a C-terminal TEV-protease cleavage site (ENLYFQS). This gene was subcloned into the pET-21a(+) expression plasmid using the NdeI and XhoI restriction sites, encoding for a C-terminal (His)_6_ affinity tag when read in-frame. The complete expression plasmid was transformed into chemically competent BL21(DE3) Δ*acrB* cells, spread onto Luria-Bertani (LB) agar plates supplemented with 100 μg/mL ampicillin, and grown overnight at 37 °C. Colonies from these plates served as the source of *E. coli* for small-scale starter cultures. *Ec*ArgRS was purified in essentially the same manner as *Sc*ATE1 with the following modification: during size-exclusion chromatography a buffer composed of 25 mmol/L Tris, pH 7.5, 100 mmol/L NaCl, 1 mmol/L TCEP, 0.35 mol/L glycerol was used. Under these conditions, *Ec*ArgRS eluted on a 120 mL gel filtration column as two species: one aggregated species and one apparent hexameric species. Only the hexameric species was pooled for downstream arginylation assays. Protein concentration was determined using the Lowry assay, and purity was assessed via SDS-PAGE analysis (acrylamide mass fraction of 15 %).

### Iron Content Determination

Iron content was determined spectrophotometrically using a modified version of the ferrozine assay.^78,79^ Briefly, protein was precipitated using 5 mol/L trichloroacetic acid (TCA). The supernatant was decanted and subsequently neutralized with saturated ammonium acetate. To this solution, excess ascorbic acid and 0.30 mmol/L ferrozine (final concentration) were added. Absorbance measurements of samples made in at least triplicate were taken at 562 nm. The concentration of Fe^2+^ was then determined assuming a Fe^2+^-ferrozine complex with an extinction coefficient (ε_562_) of ≈ 28 mmol L^−1^ cm^−1^ ^79^ (26.98 mmol L^−1^ cm^−1^ ± 0.96 mmol L^−1^ cm^−1^)^78^, and these data were corrected against residual iron present in buffer constituents.

### Anoxic Reconstitution

Apo protein samples were reconstituted in an anoxic chamber containing a N_2_/H_2_ atmosphere and operating at < 7 mg/m^3^ (5 ppm) O_2_. Briefly, protein was brought into the anoxic chamber and allowed to equilibrate with the anaerobic chamber’s atmosphere overnight at 6 °C with shaking. Protein was then diluted to 100 μmol/L in reconstitution buffer comprising 25 mmol/L 3-morpholinopropane-1-sulfonic acid (MOPS) or 25 mmol/L Tris, pH 7.5, 300 mmol/L KCl, 1 mmol/L DTT, 0.7 mol/L glycerol. 10 mmol/L stock FeCl_3_ was first titrated into the apo protein until 4 mole eq. to 10 mole eq. had been added (as warranted) with 15 min with shaking at 6 °C between the addition of each mole eq. of Fe^3+^. 10 mmol/L stock Na_2_S was then titrated into the iron-bound protein in the same manner. Afterwards, protein was allowed to equilibrate with FeCl_3_ and Na_2_S overnight at 6 °C with shaking. Particulate matter was removed by first centrifuging at 10000×*g* anoxically for 10 min, 4 °C and then by filtration through a filter with a pore size of 0.22 μm. Excess iron and sulfide were removed by buffer exchanging three times into fresh buffer. Iron and protein contents were determined as described (*vide supra*).

### Electronic Absorption (EA) and Circular Dichroism (CD) Spectroscopies

Electronic absorption (EA) spectra were recorded at room temperature on a UV-visible spectrophotometer. Samples were contained within a 1 cm UV-transparent cuvette, and data were acquired from 900 nm to 250 nm with the instrument set to a spectral bandwidth of 2 nm. Heme titrations were carried out similar to previously-described methods.^33^ Circular dichroism (CD) spectra were recorded on a nitrogen-flushed spectropolarimeter operating at room temperature. Samples were contained within a quartz cuvette with a pathlength of 0.3 cm, and data were acquired from 800 nm to 300 nm with the instrument set to a spectral bandwidth of 1 nm. The data presented are an average of 5 scans.

### X-ray Absorption Spectroscopy (XAS)

Samples containing ≈ 0.5 mmol/L to 2 mmol/L iron (final concentration) in buffer plus 3.6 mol/L ethylene glycol were aliquoted either under oxic conditions or anoxically (as warranted) into cells wrapped with tape, flash frozen in N_2(l)_ and stored at −80 °C until data collection. X-ray absorption spectra (XAS) were collected on beamlines 7-3 and 9-3 at the Stanford Synchrotron Radiation Lightsource (Menlo Park, CA) as replicates when possible. Extended X-ray absorption fine structure (EXAFS) of Fe (7210 eV) was measured using a Si 220 monochromator with crystal orientation ϕ = 90°. Samples were measured as frozen aqueous glasses at 15 K, and the X-ray absorbance was detected as Kα fluorescence using either a 100-element (beamline 9-3) or 30-element (beamline 7-3) Ge array detector. A Z-1 metal oxide filter (Mn) and slit assembly were placed in front of the detector to attenuate the elastic scatter peak. A sample-appropriate number of scans of a buffer blank were measured at the absorption edge and subtracted from the raw data to produce a flat pre-edge and eliminate residual Mn K*β* fluorescence of the metal oxide filter. Energy calibration was achieved by placing a Fe metal foil between the second and third ionization chamber. Data reduction and background subtraction were performed using EXAFSPAK.^80^ The data from each detector channel were inspected for drop outs and glitches before being included into the final average. EXAFS simulation was carried out using the program EXCURVE 9.2 as previously described.^81–83^ The quality of the fits was determined using the least-squares fitting parameter, *F*, which is defined as:

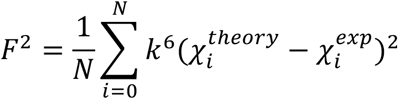

and is referred to as the fit index (FI). Low yields precluded the analysis of *Sc*ATE1 purified under anoxic conditions.

### Electron Paramagnetic Resonance (EPR) Spectroscopy

Samples containing ≈ 100 μmol/L to 600 μmol/L iron (final concentration) in buffer plus 3.6 mol/L ethylene glycol were aliquoted either aerobically or anaerobically (as warranted) into standard quartz X-band electron paramagnetic resonance (EPR) tubes with an outer diameter of 4 mm outer diameter and flash-frozen in N_2(l)_. Spectra were collected at temperatures indicated in the figure legend using a commercial EPR spectrometer system equipped with a high-sensitivity, TE-mode, CW resonator and commercial temperature-control unit. The uncertainty on the reported *g* values is 0.0005, using the manufacturer-reported field (0.08 mT) and frequency (0.00005 GHz) accuracies. The maximum, minimum and baseline-crossing points of peaks were used to determine magnetic field positions for *g* values. Calculated *g* values (from magnetic field values) agree with *g* values directly reported by the spectral analysis software provided with the commercial instrument to within 0.001.

### NMR Spectroscopy

Anoxically reconstituted samples containing *ca*. 200 μmol/L *Sc*ATE1 were exchanged into 50 mmol/L Tris, pD 7.5, 100 mmol/L KCl, 1 mmol/L DTT using a desalting column and loaded into an NMR tube with an outer diameter of 3 mm that was capped to avoid exposure to air. Paramagnetic 1D ^1^H NMR spectra were acquired at 10 °C on a 500 MHz spectrometer equipped with a room temperature probe and processed. Data collection was carried out using excitation sculpting to suppress signals from residual water. Spectra were collected with 1024 transients, 4096 time domain points, and a spectral width of 25000 Hz.

### Arginylation Assays

Arginylation activity was measured *in vitro* using a modified version of an established mass-spectrometry protocol.^63^ Stock RNase-free 4x assay buffer was prepared, comprising 200 mmol/L (4-(2-hydroxyethyl)-1-piperazineethanesulfonic acid) (HEPES), pH 7.5, 100 mmol/L KCl, and 60 mmol/L MgCl_2_. Stock DTT was added to 0.4 mmol/L final concentration in the 4x assay buffer immediately prior to setting up samples for arginylation assays. The initial reaction mixture was assembled, comprising 0.7 mg/mL *E. coli* or yeast tRNAs, 4 μmol/L *Ec*ArgRS, and 0.5 mmol/L *L-*Arg (all final concentrations) in 1x assay buffer. The initial aminoacylation reaction was initiated by the addition of a solution of Na_2_ATP to 2.5 mmol/L final concentration. This reaction mixture was incubated at 37 °C with shaking for 2 h to allow for the Arg tRNA^Arg^ to be generated. Afterwards, α-lactalbumin (the Arg acceptor protein) and ATE1 were added to 0.125 mg/mL and 2 μmol/L final concentrations, respectively. This reaction mixture was then incubated at 37 °C with shaking for 40 min to allow for the arginylation reaction to take place. The reaction was immediately halted by flash-freezing on N_2(l)_. Prior to analysis, the reaction mixture was desalted via HPLC and then immediately analyzed on a Fourier transform ion cycoltron resonance mass spectrometer. Negative controls lacking the ATE1 construct showed the expected molecular weight of unmodified α-lactalbumin (14176 g/mol (Da)) with no signs of N-terminal modification. SDS-PAGE analysis (acrylamide mass fraction of 15 %) on the final reaction mixture revealed no signs of proteolysis of either *Sc*ATE1, *Ec*ArgRS, or α-lactalbumin proteins after the reaction was completed.

### Bioinformatics

All ATE1 sequences were obtained from the Universal Protein Resource (UniProt) Knowledgebase and Reference Clusters (http://www.unprot.org) or the National Center for Biotechnology Information (http://www.ncbi.nlm.nih.gov/). Sequences were retrieved by standard protein-protein BLAST searches (blastp) using *Sc*ATE1 as an input. Conserved amino acids were identified via multiple sequence alignments that were performed using JalView v. 2.7 ^84^ implementing the ClustalW algorithm and the Blosum62 matrix ^85,86^.

### Data availability

Data are available from the corresponding author upon reasonable request.

## Notes

### Competing Interest Statement

The authors have declared no competing interest.

