## Supplemental Information for "Iron-sulfur clusters are involved in post-translational arginylation"

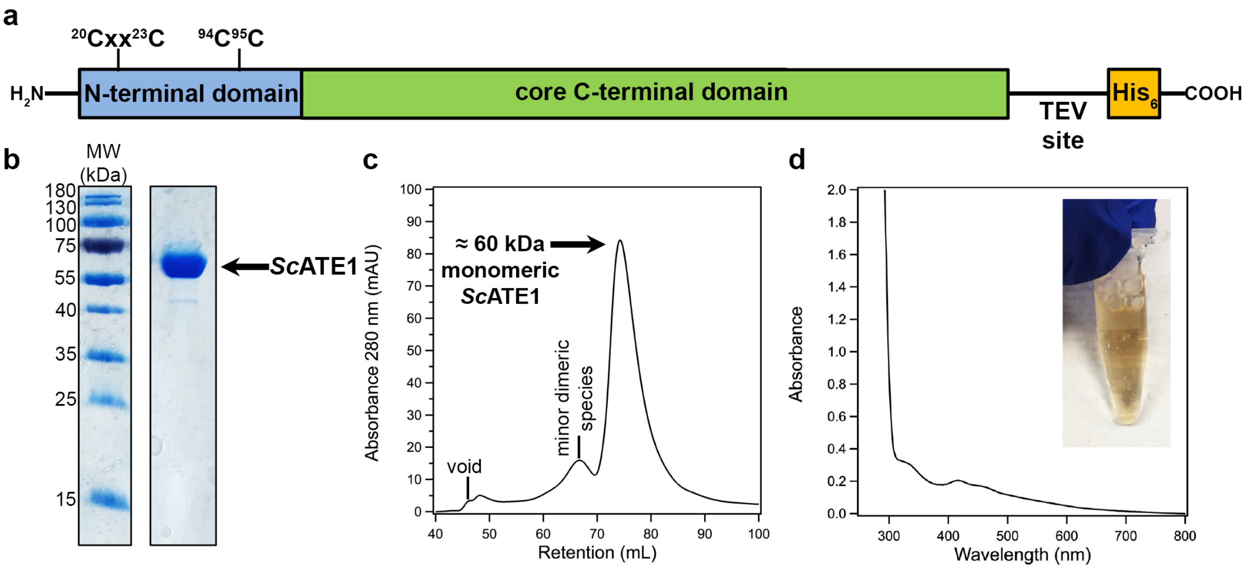


**Supplementary Figure 1**. Expression, purification, and preliminary characterization of *Saccharomyces cerevisiae* ATE1 (*Sc*ATE1). **a**. Schematic cartoon of the *Sc*ATE1 construct, with the N-terminal domain (site of [Fe-S] cluster binding) in blue, the core C-terminal domain in green, and the C-terminal ($His_6$) tag in yellow. The expression construct contains all amino acids encoded by the *ate1* gene of *Saccharomyces cerevisiae* (Uniprot identifier P16639). **b**. SDS-PAGE (acrylamide mass fraction of 15 %) analysis of *Sc*ATE1 post metal-affinity and size-exclusions chromatographies. *Sc*ATE1 has an approximate molecular weight of 60000 g/mol (60 kDa). **c**. Size-exclusion chromatogram of purified *Sc*ATE1. The major oligomeric species appears to be monomeric based on molecular-weight standards. A minor dimeric species is seen at earlier retention volumes, although this species is variable and comprises < 5 % of our best protein preparations. **d**. Concentrated *Sc*ATE1 expressed in the presence of heme precursors δ-aminolevulinic acid and ferric citrate and purified under oxic conditions displays a brown color (inset), characteristic of [Fe-S] clusters. The electronic absorption spectrum of this purified protein confirms that [Fe-S] clusters are present at low quantities under these expression and purification conditions after size-exclusion chromatography.

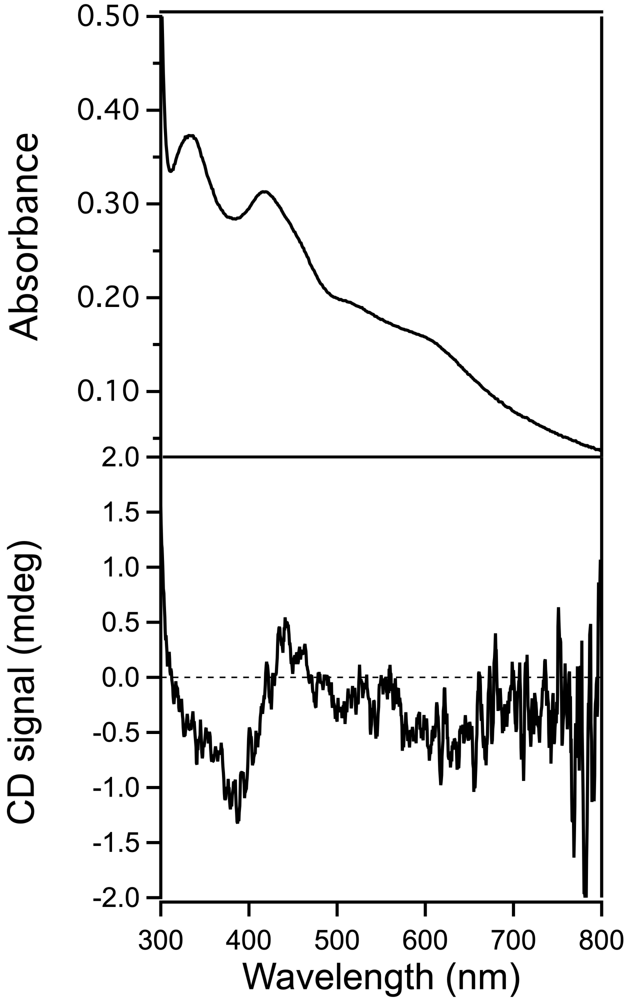


**Supplementary Figure 2**. The electronic absorption spectrum (top) and circular dichroism (CD) spectrum (bottom) of *Sc*ATE1 expressed in the presence of *L*-Cys, ferric citrate, and iron-sulfur cluster (ISC) biosynthesis machinery, and subsequently purified under oxic conditions. The spectra are characteristic of the presence of a $[2Fe\text{-}2S]^{2+}$ cluster.

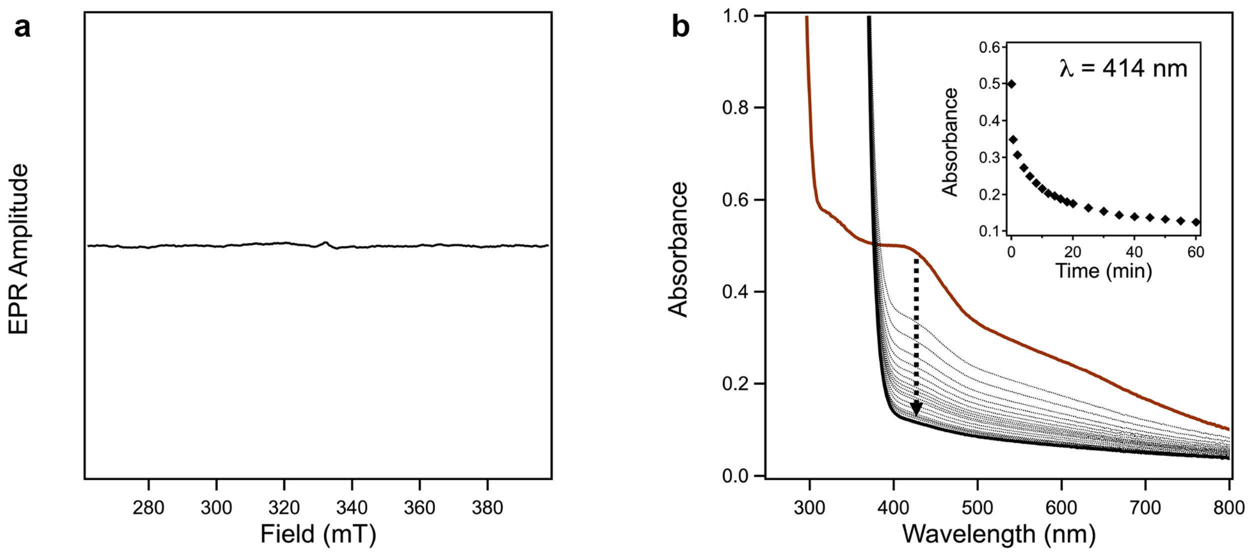


**Supplementary Figure 3**. Characterization and reactivity of the *Sc*ATE1 [2Fe-2S]$^{2+}$ cluster. **a**. The EPR spectrum of *Sc*ATE1 displays no EPR signal after purification under oxic conditions, consistent with either an antiferromagnetically-coupled [2Fe-2S]$^{2+}$ cluster (supported by other spectral assignments) or an antiferromagnetically-coupled [4Fe-4S]$^{2+}$ cluster (not supported by other spectral assignments under oxic conditions). Spectral acquisition parameters were as follows: frequency = 9.38 GHz, temperature = 20 K, modulation amplitude = 0.5 mT, modulation frequency = 100 kHz, 1024 points, conversion time = 87.89 ms, microwave power = 9.5 mW, 16 scans. **b**. Reaction of *Sc*ATE1 [2Fe-2S]$^{2+}$ with sodium dithionite under anoxic conditions results in a time-dependent bleaching (dotted) of the electronic absorption spectrum, from the starting spectrum (brown, solid) to the final spectrum (black, solid). Inset: time-dependent loss of the 414 nm feature of the electronic absorption spectrum of *Sc*ATE1 [2Fe-2S]$^{2+}$after addition of sodium dithionite.

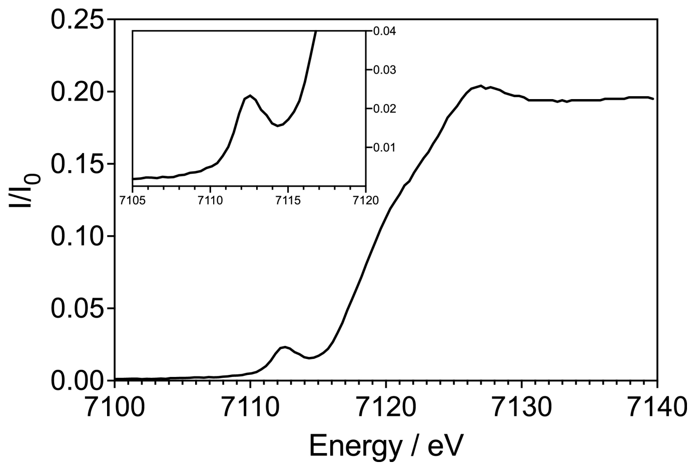


**Supplementary Figure 4**. The XANES spectrum of *Sc*ATE1 reconstituted under anoxic conditions Inset: expanded pre-edge feature of the 1s→3d transition at 7112.5 eV, consistent with tetrahedral Fe sites.

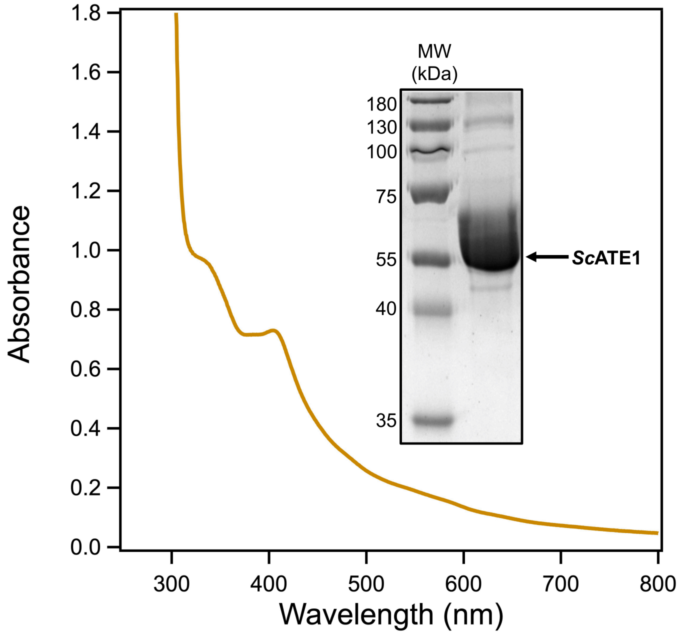


**Supplementary Figure 5**. *Sc*ATE1 expressed in anaerobic *E. coli* and purified under anoxic conditions binds a higher-order cluster. The electronic absorption spectrum (orange) displays peak maxima indicative of a mixture of $[4Fe\text{-}4S]^{2+}$ and $[3Fe\text{-}4S]^{+}$ clusters. Inset: the SDS-PAGE analysis of *Sc*ATE1 expressed in anaerobic *E. coli* and purified under anoxic conditions demonstrates good final purity, although the total protein yield is significantly diminished. Purified *Sc*ATE1 has an approximate molecular weight of 60000 g/mol (60 kDa).

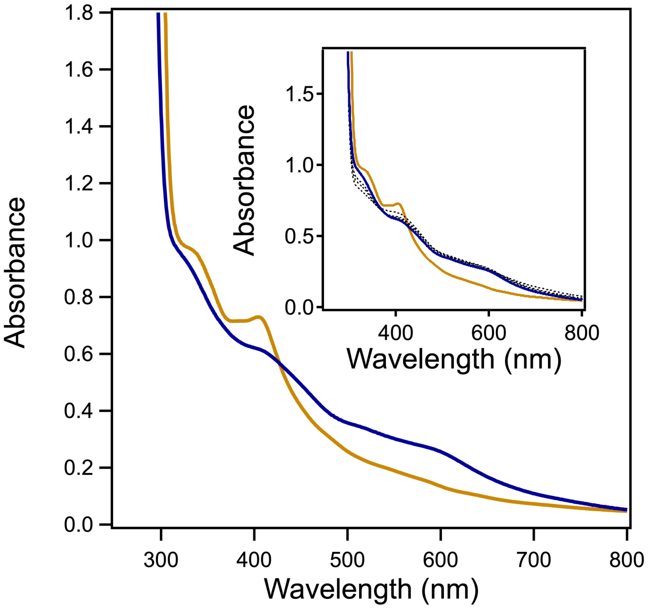


**Supplementary Figure 6**. *Sc*ATE1 expressed in anaerobic *E. coli* and purified under anoxic conditions is oxygen sensitive. The electronic absorption spectrum of *Sc*ATE1 purified anoxically before (orange) and after (blue) exposure to ambient atmosphere. Inset: time-dependent electronic absorption scans (dotted) of the oxidation of *Sc*ATE1 purified anoxically. The majority of the protein (estimated $> 90$ %) is oxidized in $< 5$ min.

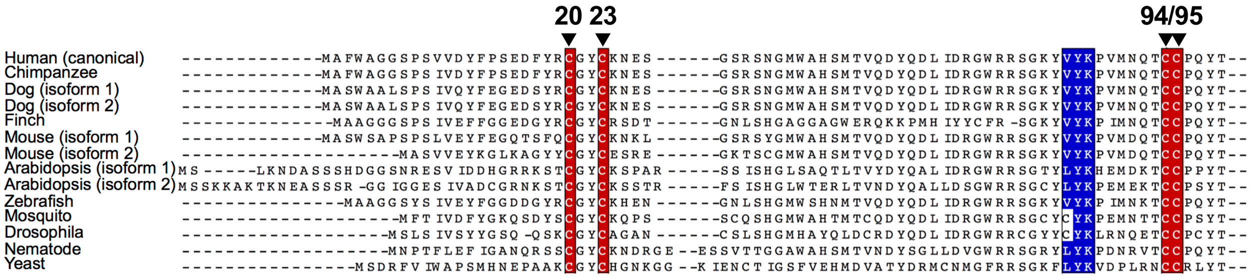


**Supplementary Figure 7**. Partial protein sequence alignments of the N-terminal domains of eukaryotic ATE1s. For brevity's sake, a small, curated set of exemplary ATE1s across the eukaryotic domain are shown: human (*Homo sapiens*; Uniprot ID: P16639), chimpanzee (*Pan troglodytes*; Uniprot ID: H2Q2P4), dog (*Canus familiaris*; isoform 1 Uniprot ID: J9P4X6; isoform 2 Uniprot ID: E2RS45), finch (*Poephila guttata*; Uniprot ID: H0ZM06), mouse (*Mus musculus*; isoform 1 Uniprot ID: Q80YP1; isoform 2 Uniprot ID: Q4FCQ6), arabidopsis (*Arabidopsis thaliana*; isoform 1 Uniprot ID: Q9ZT48; isoform 2 Uniprot ID: Q9C776), zebrafish (*Brachydanio rerio*; Uniprot ID: G8XPI0), mosquito (*Aedes aegypti*; Uniprot ID: Q178G8), drosophila (*Drosophila melanogaster*; Uniprot ID: O96539), nematode (*Caenorhabditis elegans*; Uniprot ID: P90914), and yeast (*Saccharomyces cerevisiae*; Uniprot ID: P16639). The wholly-conserved Cys residues involved in cluster binding are highlighted in red, and the strongly-conserved "LYK" motif that is involved in recognition of iron-sulfur cluster biogenesis machinery is highlighted in blue. The numbering of the Cys residues at the top of the figure is based on the yeast (*Sc*ATE1) protein sequence.

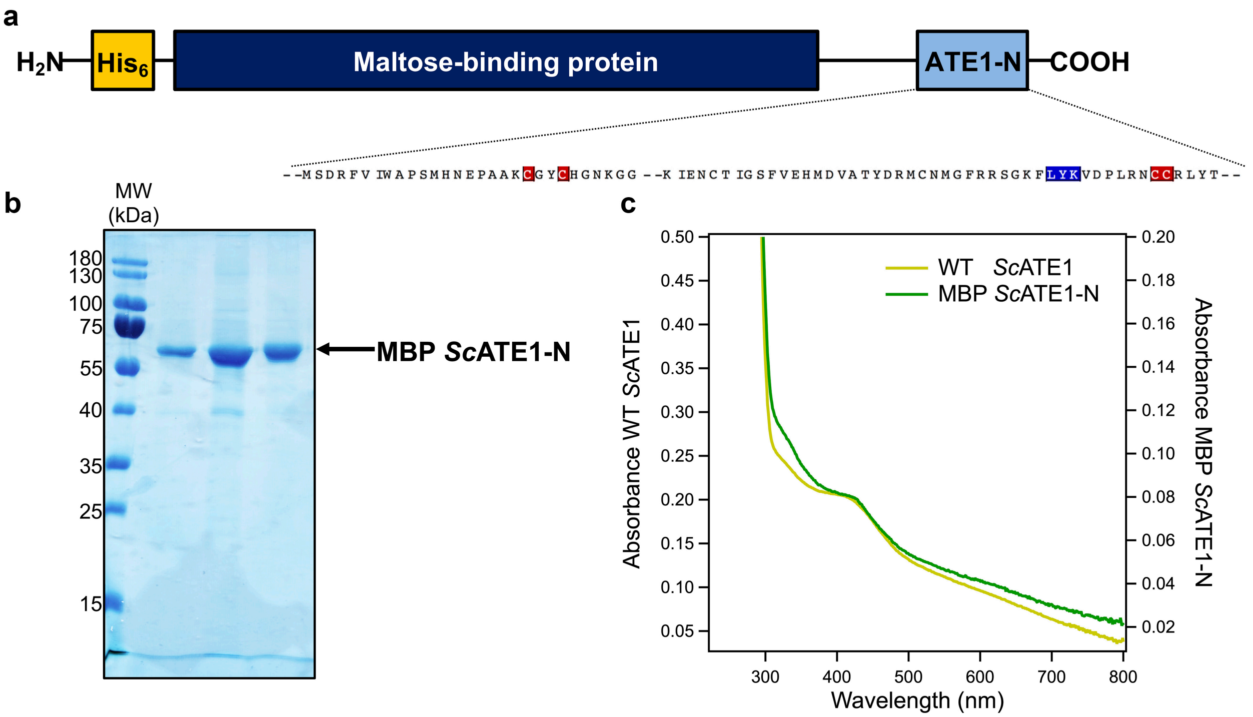


**Supplementary Figure 8**. Features of the MBP *Sc*ATE1-N construct. **a**. Schematic cartoon of the MBP *Sc*ATE1-N construct. The construct consists of an N-terminal (His)$_6$ tag (for orthogonal purification and Western blotting, yellow box), a maltose-binding protein segment for solubility (dark blue box), and the N-terminal 146 amino acids of *Sc*ATE1 (light blue box). The expansion of the light blue box approximates the locations of the conserved Cys residues (in red) and the "LYK" motif (in blue). **b**. SDS-PAGE (acrylamide mass fraction of 15 %) analysis of MBP *Sc*ATE1-N post amylose column. MBP *Sc*ATE1-N has an approximate molecular weight of 61100 g/mol (61.1 kDa). **c**. Overlay of the electronic absorption spectrum of WT *Sc*ATE1 reconstituted under anoxic conditions (yellow) and MBP *Sc*ATE1-N also reconstituted under anoxic conditions (green). The spectral are almost identical, indicating that the reconstituted $[4Fe\text{-}4S]^{2+}$ cluster is bound in the *Sc*ATE1 N-terminal domain.

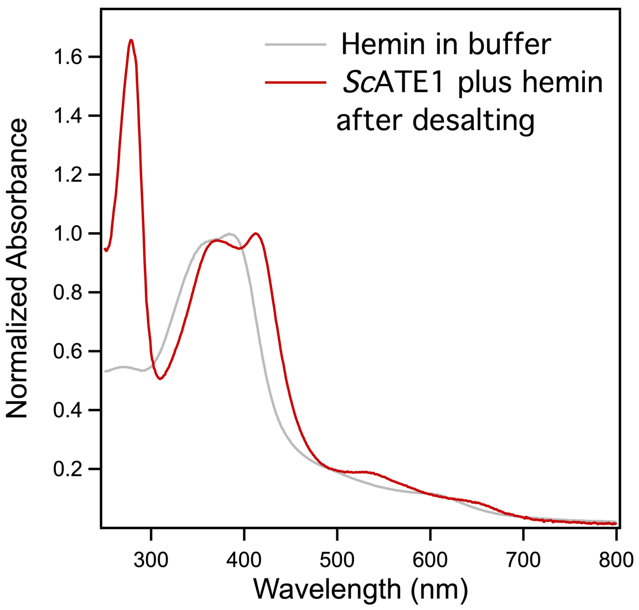


**Supplementary Figure 9**. *Sc*ATE1 with titrated hemin has an electronic absorption spectrum consistent with non-specific heme binding. The electronic absorption spectrum of heme added to *Sc*ATE1 is shown (red) after a desalting column to remove an unbound heme. For reference, hemin dissolved in *Sc*ATE1 size-exclusion buffer (see *Methods* section) is shown in gray.

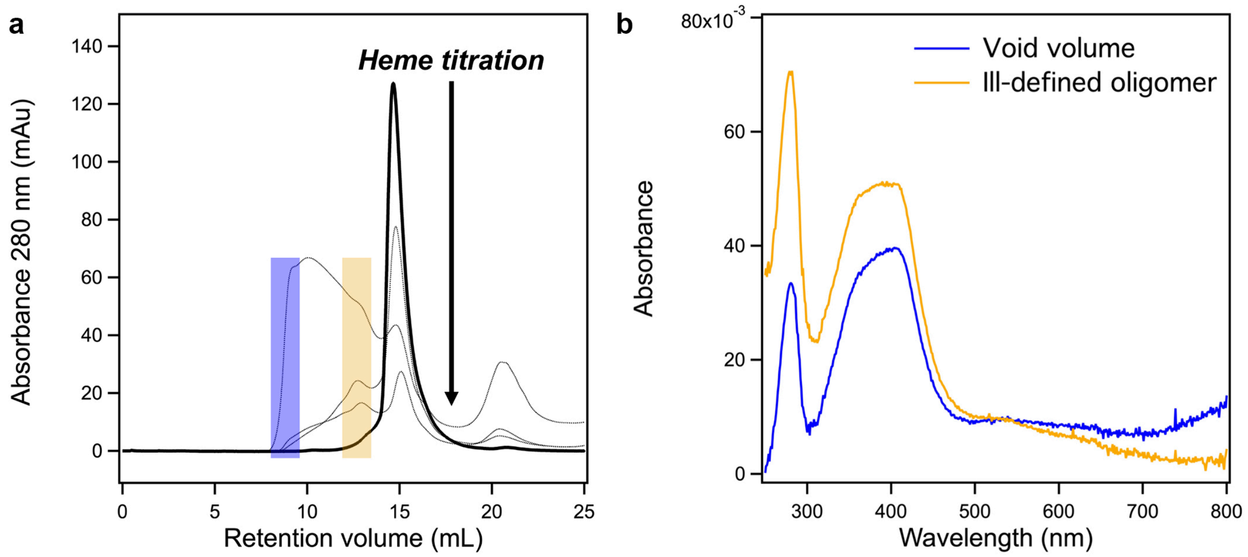


**Supplementary Figure 10**. Titration of heme into *Sc*ATE1 causes protein aggregation. **a**. In the apo form, *Sc*ATE1 is monodisperse and monomeric in solution (black solid trace) based on its retention volume (*ca*. 15 mL) compared to external standards. As heme is titrated into the protein (1, 2, 3, and 5 mole equivalents; gray dotted traces), the monodisperse and monomeric species is lost, resulting in an inactive ill-defined oligomer (*ca*. 12 mL retention volume) and inactive aggregate (*ca*. 9 mL retention volume; *i.e.*, the void volume of the column). **b**. After titration and size-exclusion chromatography, the only species that have appreciable heme bound to them are those of the inactive ill-defined oligomer (electronic absorption spectrum in orange) and the inactive aggregated species (electronic absorption spectrum in blue). Highlighted on panel **a**. are the fractions that were collected to generate the electronic absorption spectra in panel **b**.

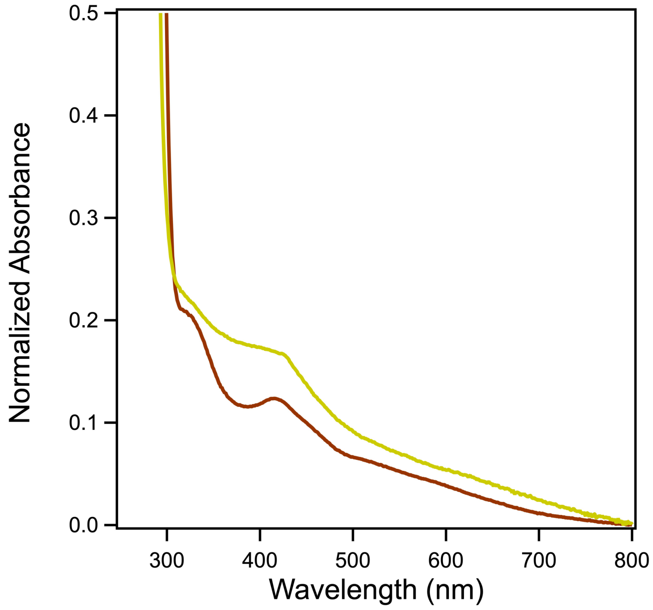


**Supplementary Figure 11**. Mouse ATE1 (*Mm*ATE1$^{1B7A}$) is also an [Fe-S] cluster-binding protein. Expression of *Mm*ATE1$^{1B7A}$ in the presence of ferric citrate and *L*-Cys and subsequent purification under oxic conditions yields an electronic absorption spectrum (brown trace) indicative of a [2Fe-2S] cluster, nearly identical to *Sc*ATE1. Apo purified *Mm*ATE1$^{1B7A}$ reconstituted under anoxic conditions yields an electronic absorption spectrum (yellow trace) indicative of a [4Fe-4S] cluster, nearly identical to *Sc*ATE1 and MBP *Sc*ATE1-N.
